# Functional wetland loss drives emerging risks to waterbird migration networks

**DOI:** 10.1101/2022.01.24.477605

**Authors:** J. Patrick Donnelly, Johnnie N. Moore, Michael L. Casazza, Shea P. Coons

**Affiliations:** Intermountain West Joint Venture - United States Fish and Wildlife Service, Migratory Bird Program, Missoula, MT, United States; Group For Quantitative Study of Snow and Ice, Department of Geosciences, University of Montana, Missoula, MT, United States; United States Geological Survey, Western Ecological Research Center, Dixon, CA, United States; Avian Science Center - University of Montana, Missoula, MT, United States

**Keywords:** agriculture, flyway, migration, shorebirds, water, waterbirds, waterfowl, wetlands, wildlife refuge

## Abstract

Migratory waterbirds (i.e., shorebirds, wading birds, and waterfowl) rely on a diffuse continental network of wetland habitats to support annual life cycle needs. Emerging threats of climate and land-use change raise new concerns over the sustainability of these habitat networks as water scarcity triggers cascading ecological effects impacting wetland habitat availability. Here we use important waterbird regions in Oregon and California, USA, as a model system to examine patterns of landscape change impacting wetland habitat networks in western North America. Wetland hydrology and flooded agricultural habitats were monitored monthly from 1988 to 2020 using satellite imagery to quantify the timing and duration of inundation—a key delimiter of habitat niche values associated with waterbird use. Trends were binned by management practice and wetland hydroperiods (semi-permanent, seasonal, and temporary) to identify differences in their climate and land-use change sensitivity. Wetland results were assessed using 33 waterbird species to detect nonlinear effects of network change across a diversity of life cycle and habitat needs. Pervasive loss of semi-permanent wetlands was an indicator of systemic functional decline. Shortened hydroperiods caused by excessive drying transitioned semi-permanent wetlands to seasonal and temporary hydrologies—a process that in part counterbalanced concurrent seasonal and temporary wetland losses. Expansion of seasonal and temporary wetlands associated with closed-basin lakes offset wetland declines on other public and private lands, including wildlife refuges. Diving ducks, black terns, and grebes exhibited the most significant risk of habitat decline due to semi-permanent wetland loss that overlapped important migration, breeding, molting, and wintering periods. Shorebirds and dabbling ducks were beneficiaries of stable agricultural practices and top-down processes of functional wetland declines that operated collectively to maintain habitat needs. Outcomes from this work provide a novel perspective of wetland ecosystem change affecting waterbirds and their migration networks. Understanding the complexity of these relationships will become increasingly important as water scarcity continues to restructure the timing and availability of wetland resources.

## 1.0 Introduction

Conservation of migratory birds is complex, requiring knowledge of species movements between distinct regions spanning hundreds to thousands of kilometers that collectively support breeding, wintering, and stopover habitats. Climate and land-use change are the major factors affecting migratory habitats and have substantially increased the risk of species declines globally (Spooner et al., 2018). Migratory birds are particularly vulnerable to these changes because of life-history strategies supported by an interdependent habitat network that can expose populations to multiple risks across their range (Zurell et al., 2018). Risks are compounded by cross-seasonal effects where environmental conditions experienced in one location (breeding grounds, wintering grounds, or stopover areas) can affect the fitness in subsequent locations leading to declines in long-term demographic performance (Sedinger and Alisauskas, 2014). While some birds have changed their migration chronology and range extent to align with shifting climate and land-use patterns (Hitch and Leberg, 2007; Visser et al., 2009), increasing environmental pressures are likely to outstrip the adaptive plasticity of many species (Schmaljohann and Both, 2017).

In arid and semi-arid mid-latitudes, migratory shorebirds, waterfowl, and wading birds, hereafter *‘waterbirds’*, rely on a limited number of important wetland areas (i.e., wetland habitat network) to connect continental movements supporting annual life-cycle events. Today, water development associated with many of these sites acts as drivers of urban development and irrigated agriculture supporting metropolitan centers and agricultural economies that account for 40% of global food production (UNESCO-UN-Water, 2020). Although growth has significantly altered most wetland and riparian ecosystems, these systems remain fundamental to biological processes sustaining migratory waterbirds. Waterbirds in some regions have adapted to landscape change by utilizing agricultural food resources and flood irrigation practices to offset historic wetland losses. (Elphick and Oring, 2003; Taft and Haig, 2005; Donnelly et al., 2021). Emerging impacts of climate change in these regions raise concerns over the sustainability of continental wetland networks as water scarcity triggers land-use change and ecological effects misaligned with waterbird habitat needs (Haig et al., 2019; Donnelly et al., 2020).

Because aridity limits wetland networks, individual sites must account for multiple ecosystem demands to support differences in species life-cycle chronology and habitat needs (sensu Roach and Griffith, 2015; Elliott et al., 2019). Climate and land-use change can disproportionately affect wetland habitats resulting in differing effects on waterbird species (Amano et al., 2020). Waterfowl in North America, for example, have benefited from proactive wetland conservation across their northern prairie breeding grounds in Canada and the United States. Although population trends of many species have increased, northern pintails *(Anas acuta)* have declined due to unforeseen impacts of shifting agricultural practices misaligned with behavioral traits of nesting hens (Podruzny et al., 2002; Duncan and Devries, 2018). Understanding the complexity of similar tradeoffs will become crucial as escalating water scarcity restructures the timing and availability of wetland habitats throughout wetland habitat networks (Kirby et al., 2008). Minimizing these risks will require a novel approach to wetland conservation that considers multi-species landscape reliance.

Wetlands in Southern Oregon and Northeast California (including the extreme northeast portion of Nevada), hereafter *SONEC*, and the Central Valley of California, USA, represent two of the most important landscapes in western North America’s waterbird migration networks (Figure 1). These regions function as interdependent landscapes in the Pacific Flyway, providing wintering, breeding, and stopover habitats that link waterbird migration from the Arctic to Central-South America (Shuford et al., 1998; Baldassarre, 2014). Collectively, the regions support habitat for over 60% of waterfowl in the western half of the continent (Petrie et al., 2013; USFWS 2020) in addition to providing essential breeding, wintering, and stopover habitats for a variety of shorebird and wading bird species (American Bird Conservancy, 2015). Both regions contain sites designated as internationally important to shorebird migration that support up to 500,000 individuals annually (Shuford et al., 1998; Senner et al., 2016). Most waterbird species move through SONEC in the fall on their way to wintering grounds in the Central Valley. Most birds have departed the Central Valley by spring and use SONEC as an important stopover site before moving north for breeding (Fleskes and Yee, 2007).

**Figure 1.**
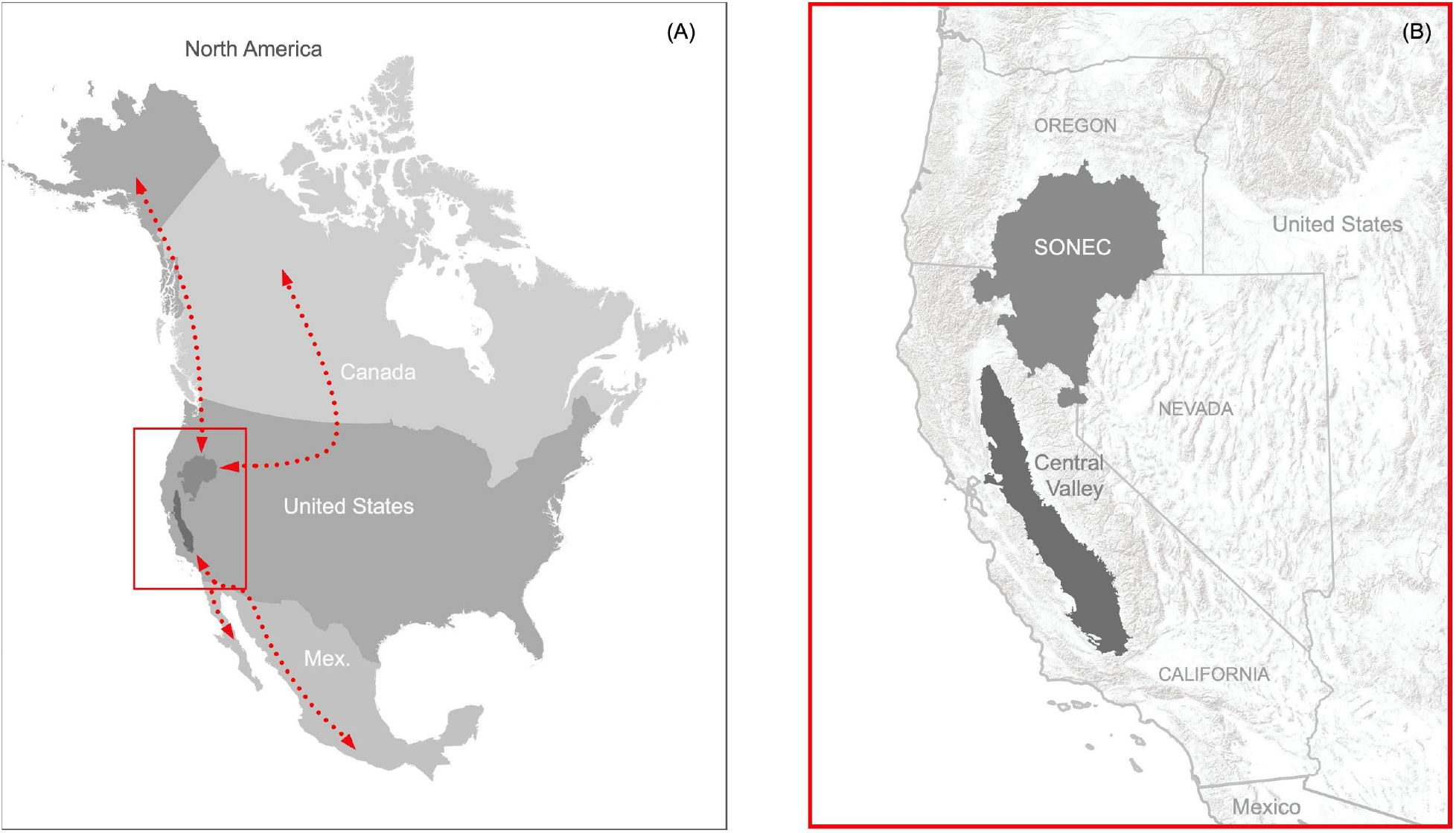
Critical landscapes connecting waterbird migration networks in western North America (A) represented by SONEC (Southern Oregon and Northeast California) and the Central Valley in the states of California, Oregon, and Nevada, USA (B).

We used SONEC and the Central Valley as a model system to identify wetland change and the resultant emerging risks to waterbird migration network in western North America. Our approach provides a unique framework to assess network risks due to a diversity of ecological and anthropogenic drivers supporting wetland functions that are distinct to each region. Results provide a novel perspective of wetland ecosystems and waterbirds that identify clear tradeoffs in potential species impacts stemming from multiple independent risks to migratory networks. Although we implement our approach using waterbird migration networks in western North America, the framework is applicable to all eight global waterbird flyways (Wetlands International, 2012), all of which are impacted by climate and land-use change (Amano et al., 2020).

## 2. Material and Methods

### 2.1 Study sites

Study sites included the SONEC and Central Valley regions in California, Nevada, and Oregon, USA (Figure 1). The SONEC region includes 11.4 million ha of the Northern Great Basin and portions of the Eastern Cascades ecoregions (Wiken et al., 2011). This area acts as a significant waterbird migration stopover site in the Pacific Flyway (Smith et al., 1989) and provides essential breeding habitat for many species, including; white-faced ibis *(Plegadis chihi)*, redheads *(Aythya americana)*, and American avocet *(Recurvirostra americana)*. Large semi-permanent wetlands also support late summer molting habitat essential to sustaining regional cinnamon teal *(Spatula cyanoptera)*, gadwall *(Mareca strepera)*, and mallard *(Anas platyrhynchos)* populations (sensu Yarris et al., 1994). Wetland freezing minimizes most waterbird use during December and January wintering periods.

The SONEC landscape is characterized by closed basins supporting palustrine emergent wetlands and littoral-lacustrine systems associated with large terminal freshwater and saline lakes, hereafter *‘closed-basin lakes’*. Most closed-basin lakes are shallow and subject to drying during prolonged periods of drought. The region is rural, with an overall human population of less than 350,000 (U.S. Census Bureau, 2021). Low-intensity farming of flood-irrigated grass hay meadows function as important agricultural resources on private lands that make up a majority of spring waterbird habitat (Donnelly et al., 2019). Other agricultural habitats include minor areas of cereal grain (e.g., wheat) that are flooded post-harvest in early spring and late fall. Public wetlands are concentrated on several large wildlife refuges managed to benefit breeding and migrating waterbirds. Climate is characterized by cold, wet winters and hot, dry summers. Wetland flooding is induced by spring runoff tied to high-elevation snowmelt. Most wetlands are flooded seasonally in late winter through early summer, after which evaporative drying reduces surface water availability. The region’s minimal reservoir storage capacity limits agriculture producers’ and public refuge managers’ ability to augment wetland water needs during drought.

The Central Valley includes 4.6 million ha of valley bottom as defined by the Central California Valley ecoregion (Wiken et al., 2011). The valley functions as one of the largest waterbird wintering areas in the Pacific Flyway. It is also recognized as a significant stopover location, connecting migrants to wintering sites in the Gulf of California, western Mexico, and Central and South America. The region provides breeding habitat for many species, including blacked-necked stilts *(Himantopus mexicanus)*, American avocets, cinnamon teal, gadwall, and mallard. Climate is characterized by temperate wet winters and hot, dry summers. Wetland conversion to industrialized agriculture beginning in the early 1900s has transformed the Central Valley into one of the most productive agricultural regions in the world, supporting 25% of U.S. food production valued at $17 billion annually (USGS 2020). Crop production is made possible through irrigation sustained by large water reclamation projects that have resulted in damming and diking of most river systems for water storage, conveyance, and flood control. Over 17 million people reside in the region, with the majority concentrated in metropolitan and urban areas embedded within the agricultural landscape (U.S. Census Bureau, 2021).

Rice cultivation makes up a majority of agricultural habitat in the Central Valley and has become crucial to sustaining wintering waterbirds (November to February) through post-harvest field flooding that decomposes leftover rice stubble (Petrie et al., 2016). Flood irrigation of rice during the growing season (May to August) can also provide important habitat for some waterbird species ((USFWS 2020)). Flooding practices associated with other crops (e.g., corn, wheat, and safflower) make up a relatively small component of available agricultural habitats (Fleskes et al., 2003). A culture of waterfowl hunting has also resulted in the substantial development of privately-owned wetlands (hereafter *duck clubs*). Most of these sites are restored agricultural lands managed for fall-winter waterfowl hunting that otherwise provide beneficial wetland habitat for waterbirds (USFWS 2020). Publicly owned wetlands are distributed across a complex of wildlife refuges managed primarily to support large concentrations of wintering waterfowl. Nearly all wetland hydrology is controlled through irrigation water conveyance and must be actively manipulated to alter the timing and duration of flooding. Exhaustive policy dictating water use combined with growing competition between agriculture, urban, and environmental demands also influences wetland hydrology and flooded agriculture patterns. High reservoir storage capacity capturing snow-melt runoff from the Sierra Nevada (mountains) allows the region to attenuate drought except during extreme conditions when water delivery supporting wetland and agricultural resources is curtailed.

### 2.2 Surface water trends

Wetland hydrology and agricultural flooding were monitored using Landsat 5 Thematic Mapper and Landsat 8 Operational Land Imager satellite imagery to depict the timing and duration of wetland surface water. Following an approach outlined by Donnelly et al. (2021), surface water conditions were measured monthly (January to December) from 1988 to 2020 as a five-year running mean beginning in 1984. Normalizing estimates in this way moderated annual climate variability influencing hydrologic conditions (Rajagopalan and Lall, 1998) and improved detectability of long-term trends. Satellite data were formatted by binning individual Landsat scenes by month and averaging results into twelve composite images for each five-year mean. Results provided 444 unique monthly measures of wetland surface water for the SONEC and Central Valley regions. The accuracy of surface water area was estimated to be 93-98% by comparison to previous work and similar methods used by Donnelly et al. (2019) that overlapped over half of our study site. The accuracy was comparable to similar time-series wetland inundation studies using Landsat data (Jin et al., 2017).

Monthly monitoring allowed individual wetlands to be separated into hydrologic regimes (hereafter ‘hydroperiods’) by totaling the monthly presence of surface water within years. Wetland totals were classified as ‘temporary’ (flooded ≤ 2 months), ‘seasonal’ (flooded > 2 and ≤ 8 months), or ‘semi-permanent’ (flooded > 8 months) using standards similar to Cowardin et al. (1979). Temporary, seasonal, and semi-permanent classes included littoral-lacustrine wetland systems associated with large closed-basin lakes found in SONEC (Cowardin et al., 1979). Wetland conditions were captured using a 30×30 meter pixel grid to account for hydrologic diversity within individual wetlands. Classification of hydroperiods provided context for wetland function important to structuring unique food resources and vegetation communities linked to waterbird foraging guilds. Flooded agriculture was omitted from the hydroperiod classification. Still, it was considered similar to seasonal and temporary wetlands for the purpose of evaluating waterbird habitat trends due to irrigation and other cultivation practices that mimicked habitat requisite of these wetland types. A description of remote sensing procedures used for wetland monitoring is provided as supplemental material (*see* Supplemental Materials - Methods, Section 1).

Wetland hydroperiod results were categorized into functional groups (Table 1) using GIS to link public-private ownership and specific ecologic and land-use characteristics to monthly surface water patterns. For example, we differentiated between natural wetlands and those actively managed through irrigation infrastructure and surface water manipulation (hereafter *managed wetlands*). To define unique functional groups, ownership was then used to subset managed wetlands by public wildlife refuges and private duck clubs. Functional group delineations were developed and stored as a polygon layer through on-screen digitizing and photo interpretation of high resolution (≤ 1 m) multispectral satellite imagery acquired after 2018. The National Agricultural Statistics Service Cropland Data Layer was used as an ancillary input to aid classification (NASS 2019). Surface water associated with large reservoirs, mining, and recreation (e.g., golf courses) was excluded due to their limited value to migratory waterbirds. Ownership was assigned using the Bureau of Land Management’s surface land ownership data (sagemap.wr.usgs.gov). Flooded agriculture occurred primarily on private lands and included minor areas on public wildlife refuges used as lure crops for wintering waterfowl.

**Table 1.** Wetland-agriculture functional groups fror Southern Oregon and Northeast California (SONEC) and the Central Valley

| Group | Description |
| --- | --- |
| Closed-basin lakes | Large terminal water bodies associated with littoral-lacustrine wetland systems in SONEC closed basins. |
| Flooded agriculture | Agricultural flooding associated with grass hay, rice, or other crop types—areas related to flood irrigation or flooding occurring post-harvest or before planting. |
| Duck clubs | Privately managed wetlands in the Central Valley maintained specifically for waterfowl hunting and wildlife— i.e., planned manipulation of surface water hydrology. |
| Private wetlands | Private un-managed or natural wetlands. |
| Wildlife refuges | Public wildlife refuges maintained specifically for wildlife through active wetland management. |
| Public wetlands | Un-managed or natural wetlands in SONEC occurring on public lands (e.g., National Forest). |

Changes to wetland hydrology in SONEC and the Central Valley were quantified by splitting monitoring results into equal periods, P1 (1988-2004) and P2 (2005-20), and measuring monthly differences using nonparametric Wilcoxon rank-order tests (Siegel, 1957). By comparing trends over long periods, we were able to minimize effects of shorter-term climate cycles (e.g. El Nino Southern Oscillation; Dettinger et al., 1998) that may have influenced results. A p-value of < 0.1 was used to represent statistical significance. Results were provided as boxplots partitioned by wetland hydroperiod (i.e., temporary, seasonal, semi-permanent) and functional groups (e.g., closed-basin lakes and cultivated rice).

Change detection analysis was used to identify wetland declines as functional or physical loss (*see* Supplemental Materials - Methods, Section 2). Functional losses were attributed to areas of diminishing surface water area (i.e., drying) associated with shifts in ecosystem water balance or water management in the absence of physical alterations to the wetland. Physical losses were attributed to land-use conversion (e.g., urban expansion or shifting agricultural practices), resulting in habitat decline. In addition, we estimated the proportional contribution of functional groups to overall wetland abundance by totaling their monthly surface water areas for P1 and P2 and dividing by their overall period sum. This approach was also used to estimate the proportional abundance of wetlands and flooded agriculture. Flooded agriculture proportions were calculated using only seasonal and temporary wetlands due to their habitat similarities supporting waterbird foraging guilds associated with shallow and seasonally intermittent surface water.

### 2.3 Waterbird habitat trends

We linked changes in monthly wetland hydrology and flooded agriculture in SONEC and the Central Valley to a suite of 33 migratory waterbirds grouped loosely by taxa and foraging guilds (Table 2). We defined an ‘other waterbird’ group that was taxonomically more diverse to act as a catch-all that included selected birds in diving, fishing, and wading guilds. Waterbird species were representative of a diversity of interdependent life-cycle events and habitat niches associated with SONEC and Central Valley. To align seasonal waterbird abundance with wetland and agricultural trends, the eBird Basic Data set (EBD) from the Cornell Laboratory of Ornithology was used (Sullivan et al., 2009). EBD was essential for constructing seasonal abundance patterns for species monitored infrequently by government wildlife agencies (e.g., shorebirds and wading birds). eBird is the largest citizen science platform globally, documenting avian-species distribution and abundance within a mobile scientific platform that ingests over 100 million observations annually.

**Table 2.** Waterbird species used in wetland cross-regional niche assessment.

| Shorebirds | Dabbling ducks |
| --- | --- |
| American avocet (*Recurvirostra americana*) | American wigeon (*Mareca americana*) |
| Black-necked stilt (*Himantopus mexicanus*) | Cinnamon teal (*Spatula cyanoptera*) |
| Dunlin (*Calidris alpina*) | Gadwall (*Mareca strepera*) |
| Greater yellowlegs (*Tringa melanoleuca*) | Green-winged teal (*Anas crecca*) |
| Lesser yellowlegs (*Tringa flavipes*) | Mallard (*Anas platyrhynchos*) |
| Long-billed dowitcher (*Limnodromus scolopaceus*) | Northern pintail (*Anas acuta*) |
| Marbled godwit (*Limosa fedoa*) | Northern shoveler (*Spatula clypeata*) |
| Western sandpiper (*Calidris mauri*) |  |
| Willet (*Tringa semipalmata*) |  |
| Wilson's phalarope (*Phalaropus tricolor*) |  |
| Wilson's snipe (*Gallinago delicata*) |  |
| Whimbrel (*Numenius phaeopus*) |  |
| Diving ducks | Other waterbirds |
| Goldeneye* (*Bucephala spp.*) | American bittern (*Botaurus lentiginosus*) |
| Bufflehead (*Bucephala albeola*) | American coot (*Fulica americana*) |
| Canvasback (*Aythya valisineria*) | Black tern (*Chlidonias niger*) |
| Scaup** (*Aythya spp.*) | Eared grebe (*Podiceps nigricollis*) |
| Redhead (*Aythya americana*) | Least bittern (*Ixobrychus exilis*) |
| Ring-necked duck (*Aythya collaris*) | Western grebe (*Aechmophorus occidentalis*) |
| Ruddy duck (*Oxyura jamaicensis*) | White-faced ibis (*Plegadis chihi*) |

\*Includes common (*B. clangula*) and Barrow's (*B. slandica*) goldeneye
\*\*Includes greater (*A. marila*) and lesser (*A. affinis*) scaup

The Auk package (Strimas-Mackey et al., 2018) was used to extract regional EBD count and presence data for all waterbird species collected from 1984 to 2020. Due to the relatively recent deployment of eBird, most observations used in our analysis were acquired post-2008. Following Strimas-Mackey (2018) EBD best practices, we restricted data to 1) standard ‘traveling’ and ‘stationary’ count protocols, 2) complete checklists, 3) observation length < 5 hours, 4) effort-distance to ≤ 5 km, and 5) number of observers ≤ 10. Results from EBD queries were binned by month (to align with wetland-agricultural monitoring outputs) and summed across years to calculate proportional waterbird abundance as a relative measure of regional bird use over time. Results were presented as bubble plots for each species by region to illustrate monthly patterns of cross-seasonal reliance (*see example*, Figure 2). Although we recognize differences in climate, weather, and disturbance can influence seasonal bird abundance, we intended to estimate long-term norms for comparison to wetland trends.

**Figure 2.**
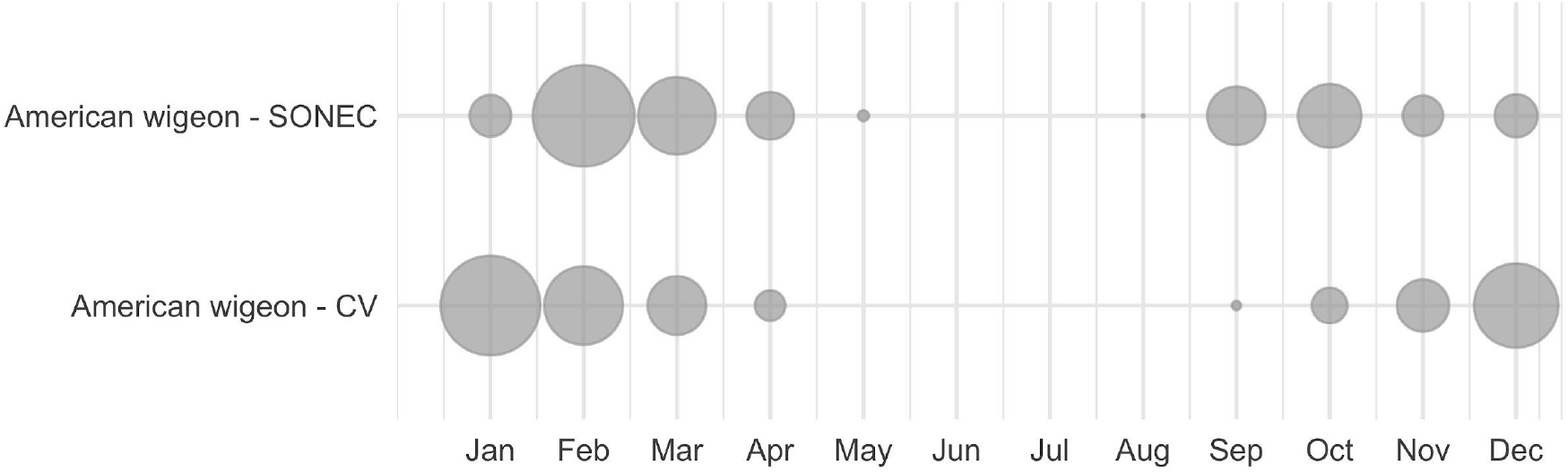
An example of Southern Oregon and Northeast California (SONEC) and Central Valley (CV) cross-seasonal waterbird distributions depicted with American wigeon. Dot size illustrates proportional abundance by region and month (Jan-Dec). High winter use (Jan) in CV shifts to SONEC during spring migration (Feb-Mar), while high SONEC use during fall migration (Sep-Nov) transitions back to Central Valley for winter (Dec). Bird absence from May to August indicates breeding is focused outside these regions.

When applied at broad scales, past studies have shown EBD observations equivalent to traditional survey efforts (Callaghan and Gawlik, 2015; Walker and Taylor, 2017). For added assurances, we compared (using non-parametric Wilcoxon tests) EBD-derived abundance distributions to results from aerial and ground surveys conducted in SONEC and the Central Valley. Although the majority of EBD observations included in our analysis were acquired post-2008, comparisons to traditional long-term (1984-2016) and near-term (2011-2017) waterbird surveys showed no significant differences (p-value <0.05) in seasonal abundance patterns (Figure S1-3). Detailed methods and results outlining this analysis are provided as supplemental material (*see* Supplemental Materials - Methods, Section 3). Patterns of seasonal waterbird abundance were linked to monthly wetland trends using a rule-based approach to identify emerging risks to niche availability broadly. Species were first assigned to one or more wetland hydroperiod classes (temporary, seasonal, and/or semi-permanent) representative of their seasonal habitat utilization (Sibley et al., 2001; Baldassarre, 2014). Flooded agriculture was an additional factor for species reliant on those habitats. Diving ducks, American coot, black tern, eared grebe, and western grebe were associated with semi-permanent wetlands that are representative of deeper open-water refugia and food resources preferred by these species. Dabbling ducks, American bittern, and white-faced ibis were associated with all wetland hydroperiod classes and flooded agriculture to encompass the diversity of their habitat utilization. As an exception, cinnamon teal, gadwall, and mallard were associated with semi-permanent wetlands from April to September when regional populations are heavily reliant on these habitats during brood rearing (Apr-Jul) and 25-40 day flightless molt periods (Aug-Sep; Kohl et al. in press). A similar rule was applied to American wigeon, green-winged teal, northern pintail, and northern shoveler to account for their minor breeding and molting occurrences in SONEC. However, April and September were excluded to prevent overlap with migrating populations that occurred in much higher abundance during those months.

Shorebird habitat assessments in SONEC were restricted to large terminal lake basins (Abert, Alkali, Goose, Harney, Honey, Summer, and Warner) identified as regionally and internationally important to sustaining populations (Senner et al., 2016). However, we acknowledge shorebird use in other wetland systems. Habitat associations included semi-permanent, seasonal, and temporary wetlands. Seasonal and temporary wetlands are commonly correlated with shallow water that are important foraging requisites for shorebird species, while semi-permanent (i.e., littoral-lacustrine) wetland trends have been identified as a key indicator of lake salinity linked trophic function supporting shorebird energetic needs (Senner et al., 2018). Shorebirds in the Central Valley were associated with all wetland classes in addition to flooded agriculture to account for a greater diversity of hydrologic conditions and habitat use driven by human-controlled flooding (Reiter et al., 2015).

Species-wetland associations were used as a template to interpret how wetland-agricultural trends were likely to affect habitat availability. To illustrate regional relationships between monthly waterbird abundance and wetland-agricultural change, species bubble plots were color-coded (Figure 3). Red (significant impacts) indicated declines to half or more of wetland types, including agriculture, supporting a species habitat niche. Yellow (moderate impact) indicated declines to a minority of associated wetland-agricultural classes. Blue (stable) indicated stable-to-increasing wetland-agricultural conditions across all associated classes. Wetland declines were determined through statistical inference using p-values < 0.1 derived from Wilcoxon rank order test (*see Methods section 2*.*2 Wetland trends*). Habitat conditions for species associated with fewer than three wetland classes (i.e., diving ducks and SONEC shorebirds) could only be assessed as ‘significantly declining’ or ‘stable/increasing’.

**Figure 3.**
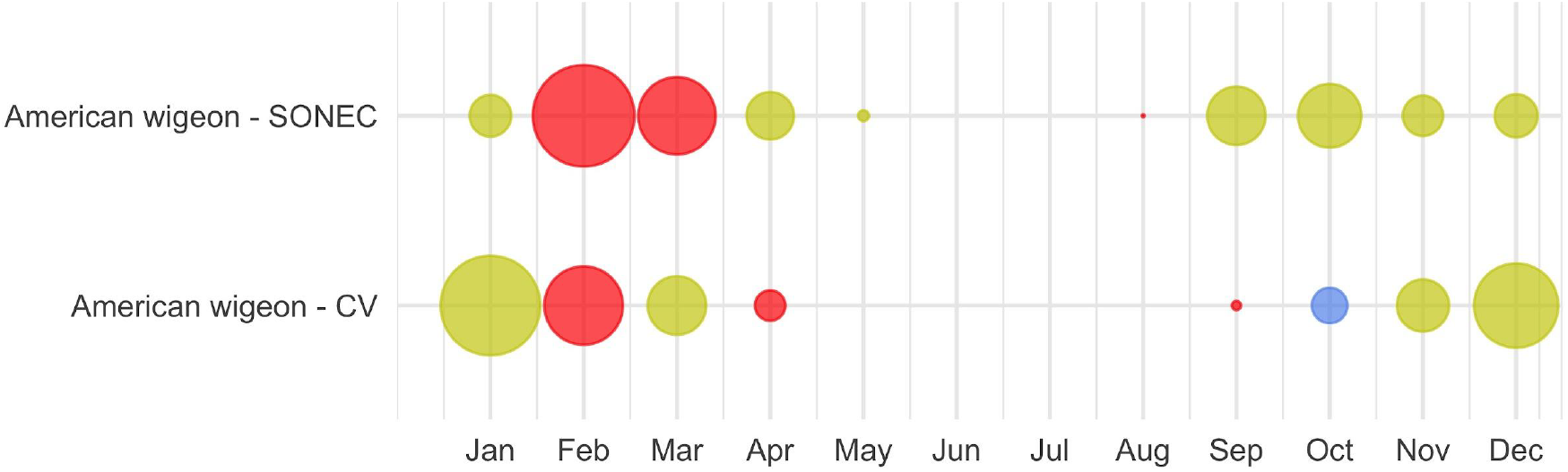
An example of Southern Oregon and Northeast California (SONEC) and the Central Valley (CV) cross-seasonal waterbird distributions depicted for American wigeon. Dot size illustrates proportional abundance by region and month (Jan-Dec). Colors represent wetland-agriculture trends underlying a species habitat niche. Red indicates ‘significant impacts’—declines to a majority of wetland-agricultural habitats utilized by a species. Yellow indicates ‘moderate impacts’—declines to a minority of wetland-agricultural habitats used. Blue indicates stable conditions.

### 2.4 Data Processing

All image processing and raster-based analyses were conducted using the Google Earth Engine cloud-based geospatial processing platform (Gorelick et al., 2017). GIS analyses were performed using QGIS (QGIS Development Team, 2020). Plotting and statistical analyses were conducted using the R environment (R Core Team, 2019; RStudio Team, 2019), including R-package tidyverse (Wickham et al., 2019).

## 3. Results

All wetland and agricultural results are provided as median differences of monthly surface water extent between P1 (1988-2004) and P2 (2005-20). Annual variability is presented using boxplots for visual comparison of monthly P1 and P2 wetland trends. Detailed results supporting our analyses are provided as supplemental material for all wetland hydroperiods and functional groups discussed below (*see* Supplemental Materials - Results, Tables S1-10, Figures. S1-7).

### 3.1 SONEC

Wetland change in SONEC was driven by functional decline as indicated by the continuous drying of semi-permanent wetlands consistent across functional groups (i.e., wildlife refuges and public-private lands). Outside periods of winter freezing, overall losses ranged from 27% (Mar) to 46% (Oct, Figure 4, Table S1). Significant seasonal and temporary wetland losses were limited to July when surface water declined 28% and 49% (Figure 4, Table S1). Compared to overall trends, seasonal wetland loss was more expansive on wildlife refuges and public lands (e.g., National Forest), showing declines beginning in May and lasting through September (Tables S3-4, Figures S6-7). Closed-basin lakes were the only functional group to exhibit positive seasonal (167%, Mar) and temporary (268%, Jun) wetland trends (Table S2, Figure S5) that offset drying in other functional groups. Flooded agriculture remained relatively stable over time, except for February and July, when surface water area declined 21% and 22% (Figure 5, Table S6). Land-use change in SONEC resulted in less than 300 ha of surface water loss in flooded agriculture, attributed to the conversion of flood irrigation to sprinkler use in grass hay agriculture.

**Figure 4.**
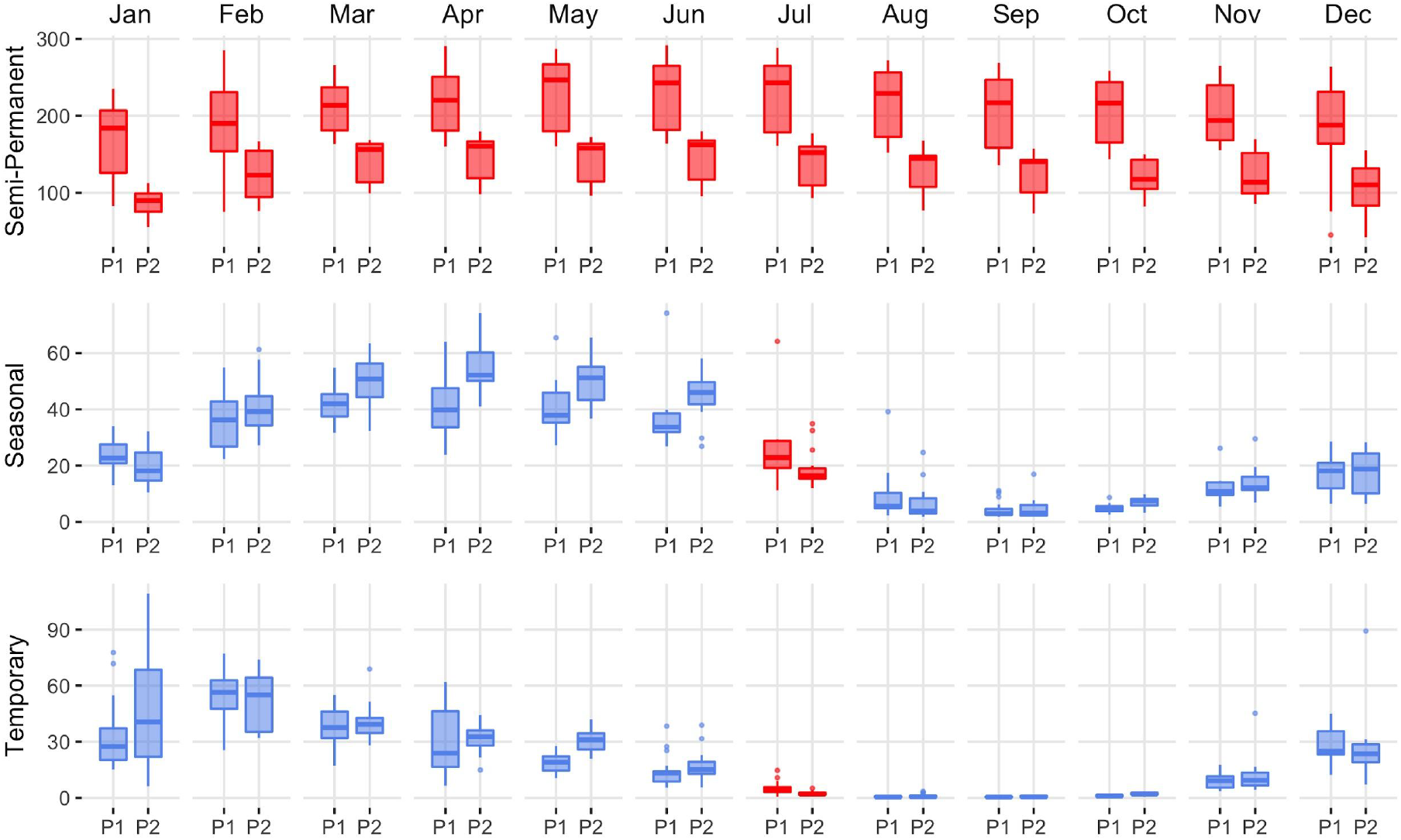
Southern Oregon and Northeast California (SONEC) overall distribution of monthly wetland abundance (kha) between 1988-2004 (P1) and 2005-20 (P2) periods. Summaries include all wetlands associated with closed basin lakes, wildlife refuges, and public-private lands. Statistical inference was determined as p-values < 0.1 derived from Wilcoxon ranked order test. Red indicates significant wetland decline, and blue indicates stable to increasing wetland abundance. Results are partitioned by wetland hydroperiod (semi-permanent, seasonal, temporary). Boxes, interquartile range (IQR); line dividing the box horizontally, median value; whiskers, 1.5 times the IQR; points, outliers.

**Figure 5.**
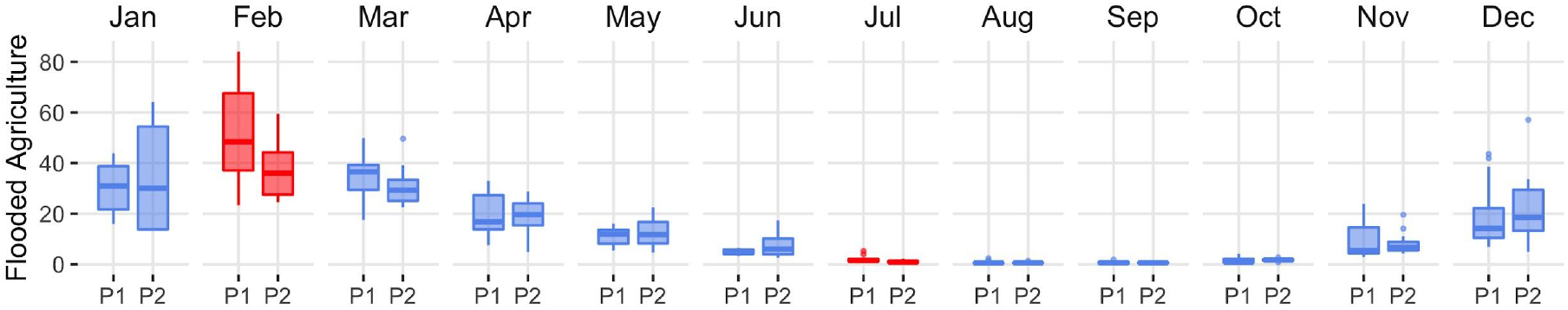
Southern Oregon and Northeast California (SONEC) distribution of monthly flooded agriculture abundance (kha) between 1988-2004 (P1) and 2005-20 (P2) periods. Statistical inference was determined as p-values < 0.1 derived from Wilcoxon ranked order test. Red indicates significant wetland decline, and blue indicates stable to increasing wetland abundance. Boxes, interquartile range (IQR); line dividing the box horizontally, median value; whiskers, 1.5 times the IQR; points, outliers.

Flooded agriculture in SONEC accounted for 76% and 73% of potential waterbird habitat annually during P1 and P2— as estimated using only seasonal and temporary wetlands due to similarities supporting waterbird guilds associated with shallow-water habitats (e.g., dabbling ducks, shorebirds, and white-faced ibis). We acknowledge, however, that this measure was based only on surface water area and did not consider greater diversity and ecological value typically attributed to wetland systems. Closed basin lakes made up the largest semi-permanent wetlands proportion, accounting for ∼76% of overall abundance (Table 3). However, most of this area was represented by open water lacustrine systems with limited habitat values for most waterbird species. Seasonal and temporary wetlands were well distributed among functional groups that made up a minimum of 21% and a maximum of 32% of overall abundance (Table 3). Wetland distributions remained relatively stable between periods, except for littoral seasonal and temporary wetlands in closed basin lakes. These increased proportionally from 32% to 43% and from 20% to 33%.

**Table 3.**
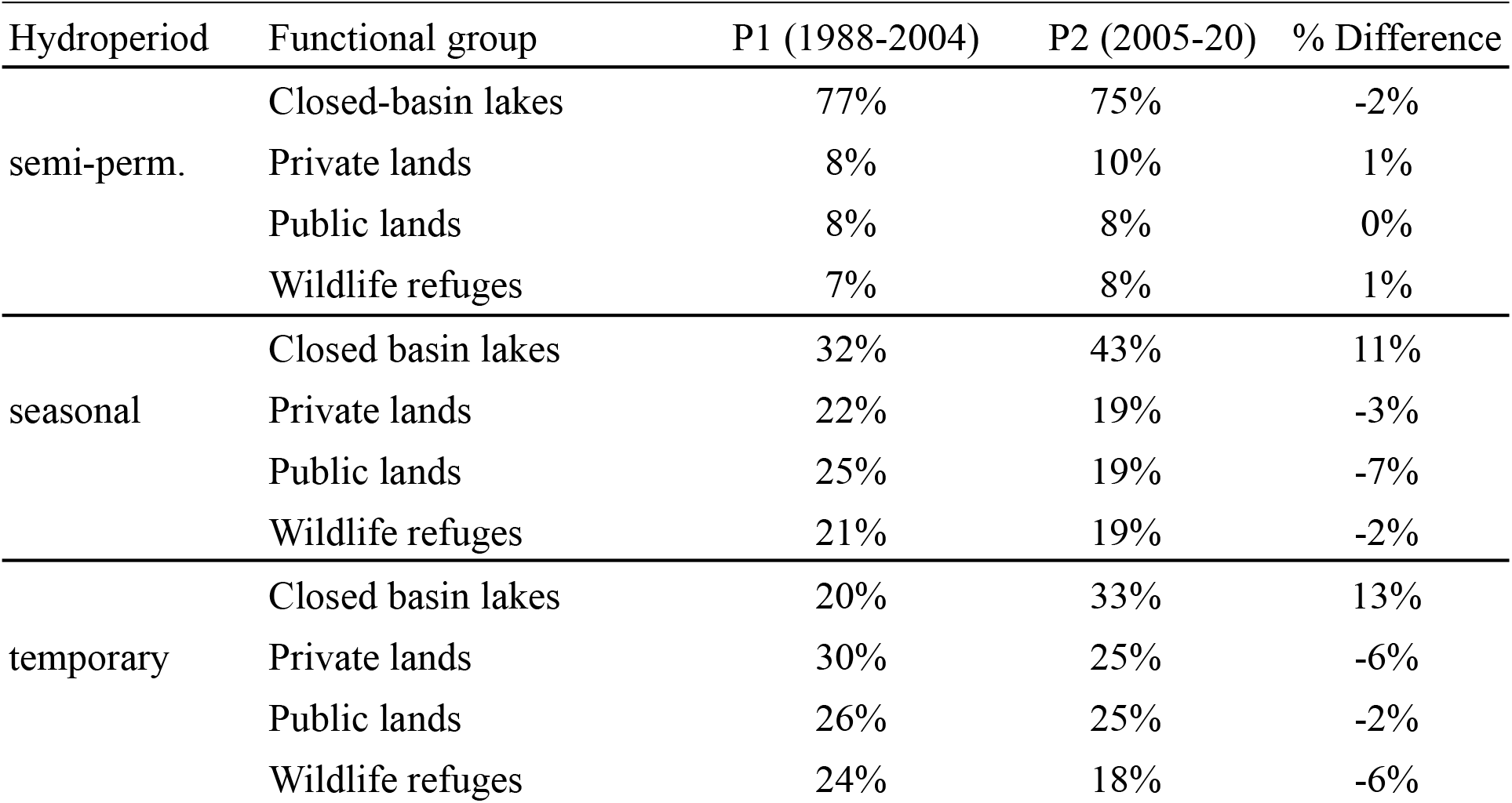
Southern Oregon and Northeast California (SONEC) proportional wetland abundance by functional group and hydroperiod for P1 (1988-2020) and P2 (2005-20).

### 3.2 Central Valley

Functional loss was the driver of wetland declines in the Central Valley as there was little evidence of physical impacts from land-use change. Drying of semi-permanent wetlands was persistent, occurring 6 out of 12 months with losses ranging from 9% (Apr) to 20% (Jan; Figure. 6, Table S7). Semi-permanent losses on wildlife refuges and duck clubs accounted for 60% and 40% of overall declines (Tables S8-9, Figures S9-10). September and October were the only months to exhibit stable semi-permanent wetland trends. Drying of seasonal and temporary wetlands was significant from April through August and September, with declines ranging from 25% to 57% (Figure 6, Table S7). Although the relative change in wetland area was low, declines coincided with annual minimums when most wetlands in the region were dry. Overall seasonal and temporary declines were representative of wetland losses on wildlife refuges and duck clubs. Other monthly declines included temporary wetlands in February (55%). Flooded agriculture increased in November, December, and January by 76%, 68%, and 29%, respectively (Figure 7, Table S10). Other monthly increases to flooded agriculture occurred in June (17%).

**Figure 6.**
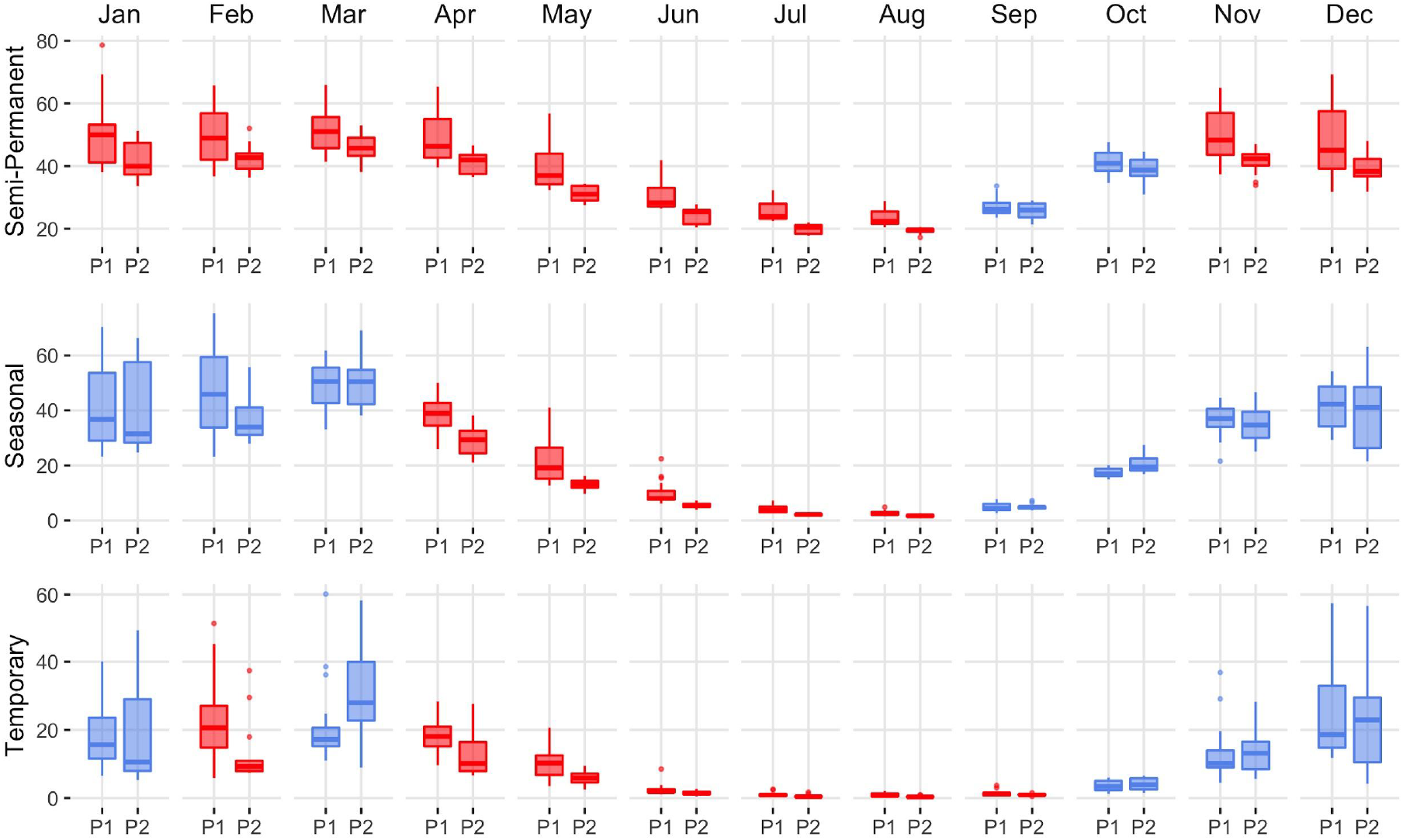
Central Valley distribution of monthly wetland abundance (kha) from 1988-2004 (P1) and 2005-20 (P2). The summary includes all wetlands on duck clubs and wildlife refuges. Statistical inference was determined as p-values < 0.1 derived from Wilcoxon ranked order test. Red indicates significant wetland decline, and blue indicates stable to increasing wetland abundance. Results are partitioned by wetland hydroperiod (semi-permanent, seasonal, temporary). Boxes, interquartile range (IQR); line dividing the box horizontally, median value; whiskers, 1.5 times the IQR; points, potential outliers.

**Figure 7.**
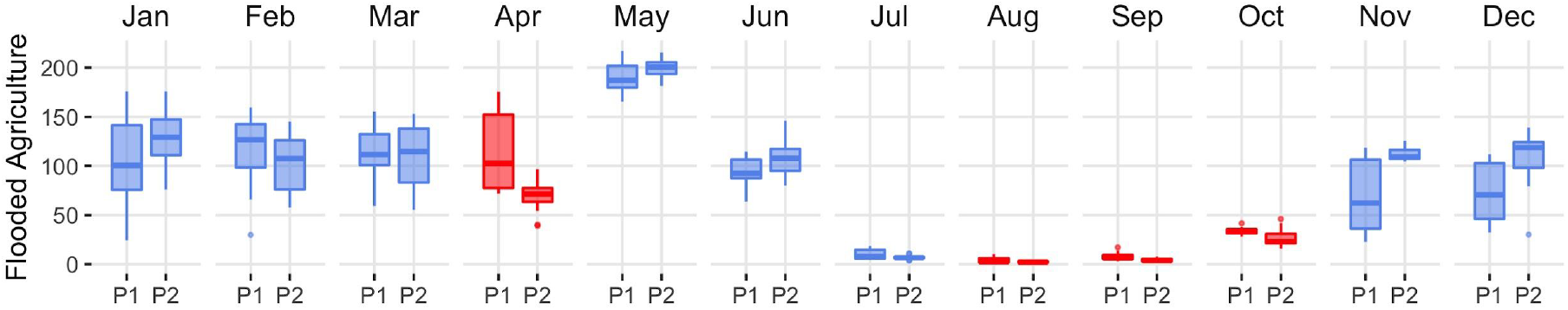
Central Valley distribution of monthly flooded agricultural abundance (kha) from 1988-2004 (P1) and 2005-20 (P2). Statistical inference was determined as p-values < 0.1 derived from Wilcoxon ranked order test. Red indicates significant decline, and blue indicates stable to expanding flooded agriculture. Boxes, interquartile range (IQR); line dividing the box horizontally, median value; whiskers, 1.5 times the IQR; points, potential outliers. Trends excluded closed basin lakes to prevent bias from large deepwater areas with minimal waterbird value.

Duck clubs accounted for over two-thirds of semi-permanent wetlands and nearly three-quarters of seasonal and temporary wetlands in the Central Valley annually, with the remainder occurring on wildlife refuges (Table 4). The proportional abundance of wetlands between duck clubs and wildlife refuges changed little over time (+/- 0.5%). Flooded agriculture made up 81% and 83% of potential waterbird habitat annually during P1 and P2. Estimates were made using only seasonal and temporary wetlands due to habitat similarities supporting waterbird foraging guilds associated with shallow and seasonally intermittent surface water. Flood irrigation of rice from April to August and post-harvest flooding for rice stubble from October to February made up the vast majority of agricultural habitat. Rice was the only waterbird habitat impacted by land-use change (i.e., physical loss) resulting from conversion to orchards and urban development. Losses were minor, representing < 4% of the cultivated footprint. Monthly patterns of flooded rice depicted by our models (Figure 7) aligned with seasonal irrigation practices (University California Davis, 2018) and estimates of the cultivated area reported for the region (Geisseler and Horwath, 2016). We acknowledge low seasonal wetland estimates in July and August were likely due to dense emergent rice cover visually obscuring areas of shallow surface water beneath.

**Table 4.** Central Valley proportional wetland abundance by functional group and hydroperiod for P1 (1988-2020) and P2 (2005-20).

| Hydroperiod | Functional group | P1 (1988-2004) | P2 (2005-20) | % Difference |
| --- | --- | --- | --- | --- |
| semi-perm. | Duck clubs | 68% | 68% | -0.5% |
|  | Wildlife refuges | 32% | 32% | 0.5% |
| seasonal | Duck clubs | 72% | 72% | 0% |
|  | Wildlife refuges | 28% | 28% | 0% |
| temporary | Duck clubs | 72% | 71% | -0.5% |
|  | Wildlife refuges | 28% | 29% | 0.5% |

### 3.3 Waterbird and wetland indicators

Wetland declines aligned with key cross-seasonal habitat needs supporting waterbirds in SONEC and the Central Valley. Indicators of significant and moderate habitat impacts were prevalent across all 33 waterbird species (Figures 8, 9). Diving ducks exhibited the broadest indications of habitat loss in SONEC and the Central Valley, resulted from semi-permanent wetland declines overlapping important stopover, breeding, molting, and wintering periods (Figure 8). Stable to increasing semi-permanent wetland trends during September and October showed only minor overlap with resident diving duck populations (i.e., ruddy duck and redhead) in the Central Valley. Similar impacts were associated with American coot, black tern, eared grebe, and western grebe because of their heavy reliance on semi-permanent wetland habitats (Figure. 9).

**Figure 8.**
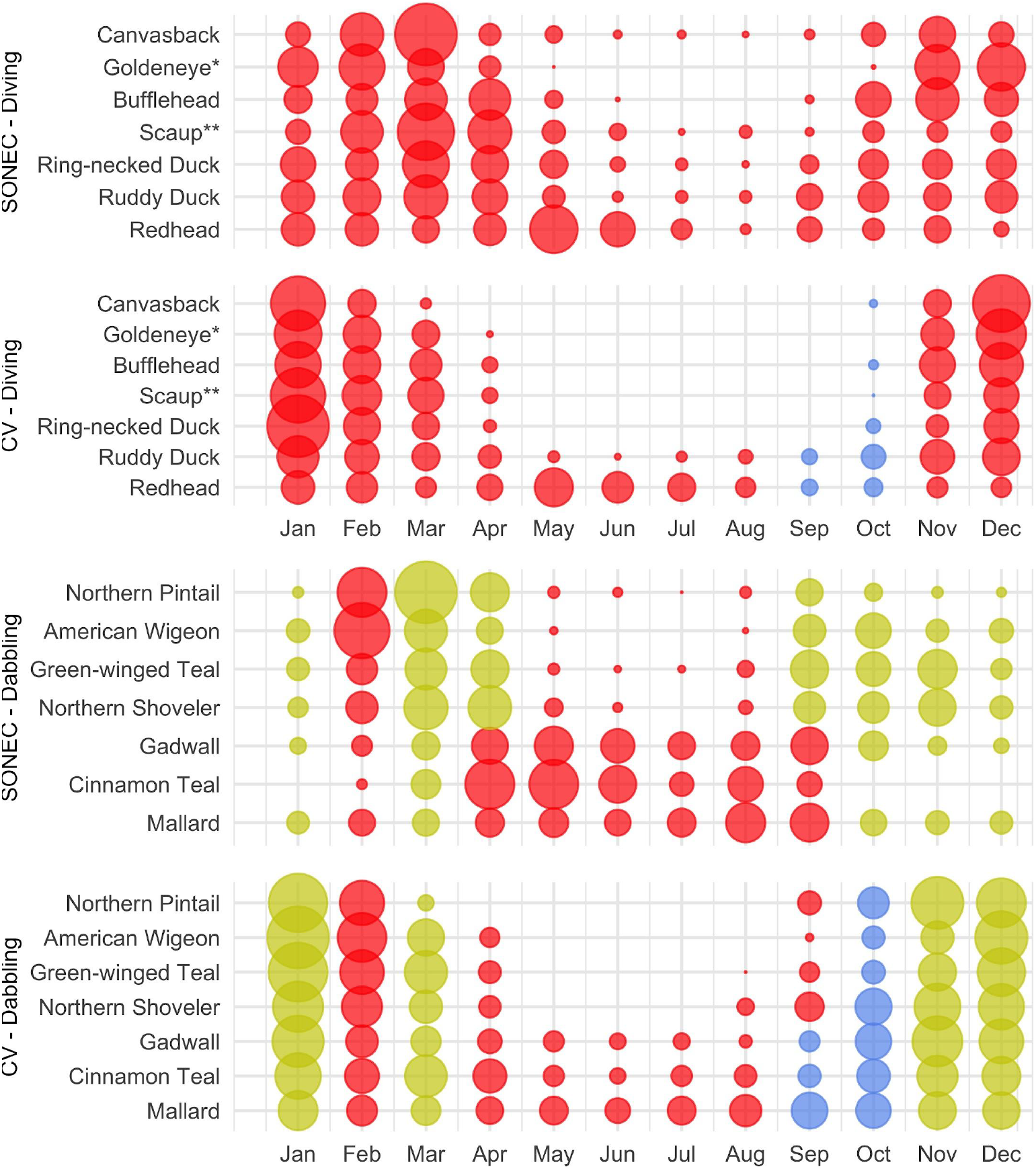
Southern Oregon and Northeast California (SONEC) and Central Valley (CV) monthly diving and dabbling duck distributions. Dot size illustrates proportional abundance from January to December. Large dots represent seasonal concentrations of birds associated with wintering and migrating behaviors. Similar-sized dots occurring over many months represent continuous bird abundance related to regional populations. Colors are indicators of habitat impacts related to changes to flooded agriculture and wetland (i.e., semi-permanent, seasonal, and temporary) abundance. Red indicates ‘significant impacts’—declines to a majority of wetland-agricultural habitats utilized by a species. Yellow indicates ‘moderate impacts’—declines to a minority of wetland-agricultural habitats used. Blue indicates stable conditions. *\*Includes common and Barrow’s goldeneye. **Includes greater and lesser scaup*.

**Figure 9.**
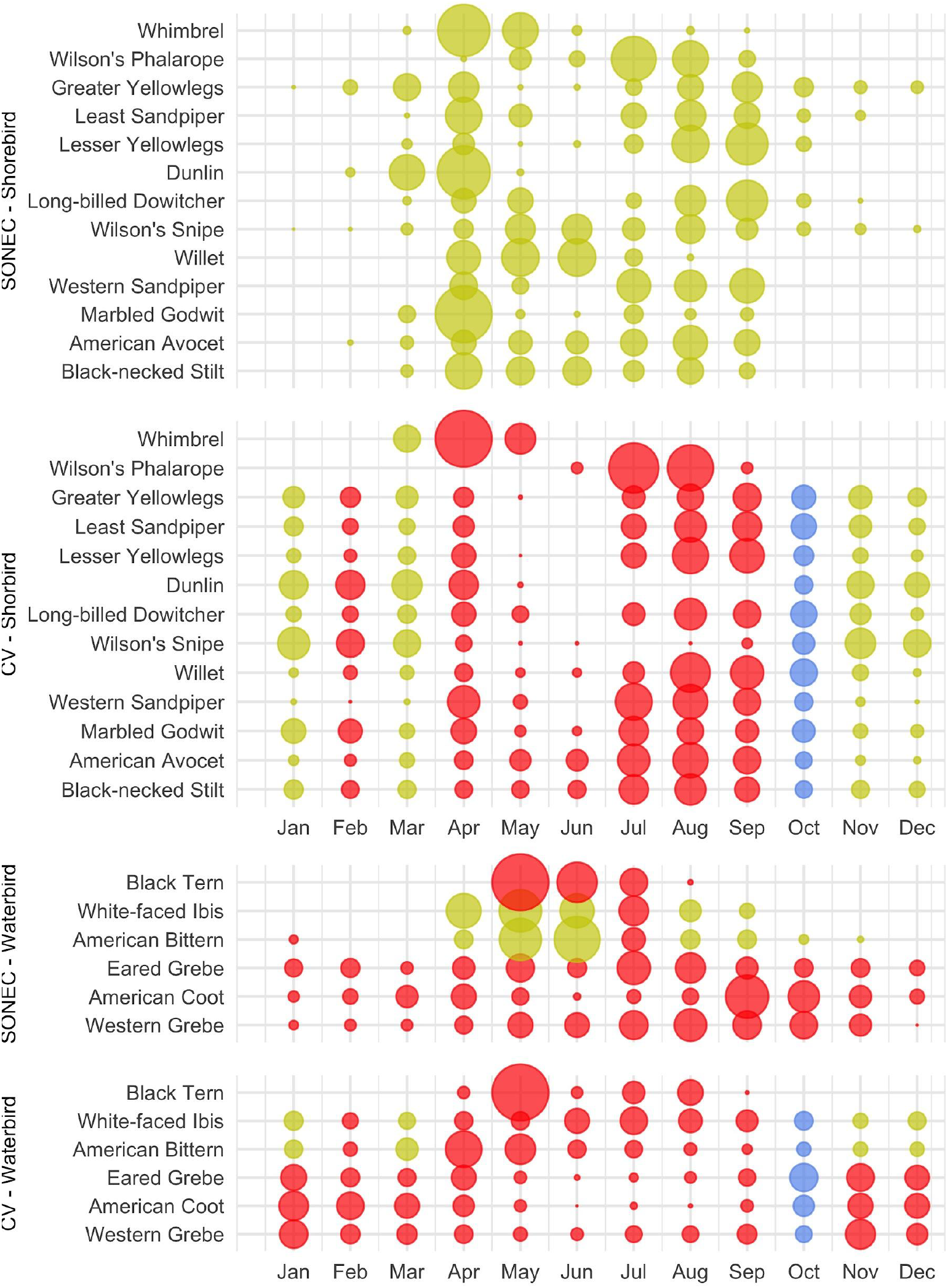
Southern Oregon and Northeast California (SONEC) and Central Valley (CV) seasonal shorebird and waterbird distributions. Dot size illustrates proportional abundance from January to December. Large dots represent seasonal concentrations of birds associated with wintering and migrating behaviors. Similar-sized dots occurring over many months represent continuous bird abundance related to regional populations. Colors are indicators of habitat impacts related to changes to flooded agriculture and wetland (i.e., semi-permanent, seasonal, and temporary) abundance. Red indicates ‘significant impacts’—declines to a majority of wetland-agricultural habitats utilized by a species. Yellow indicates ‘moderate impacts’—declines to a minority of wetland-agricultural habitats used. Blue indicates stable conditions.

Indicators of habitat declines were moderate for wintering (Dec-Jan) dabbling ducks in the Central Valley. Moderate impacts were associated with semi-permanent wetland declines on duck clubs and wildlife refuges (Figure. 8). Expansion of flooded agriculture (i.e., post-harvest flooding of rice) was also prevalent during Central Valley wintering periods (Nov-Jan), substantially increasing habitat availability. Decreasing semi-permanent and temporary wetland abundance were indicators of significant and moderate impacts to spring dabbling duck migration (Feb-Apr) in SONEC and the Central Valley. Flooded agriculture also declined 15% during February Central Valley spring migration (Table S10) but did not meet our threshold of statistical inference for wetland change—this decline resulted in a substantial loss of wetland habitat.

Habitat declines during fall dabbling duck migration were moderate for non-molting species in SONEC (Sep-Oct) and moderate and stable for all species in the Central Valley (Oct-Nov). Semi-permanent and seasonal wetland declines were the primary indicators of habitat impact. Declining semi-permanent wetlands overlapping cinnamon teal, gadwall, and mallard use were significant indicators of reduced breeding and molting habitat availability from April to September. In September, stable semi-permanent wetland trends showed only minor overlap with dabbling duck molt periods in the Central Valley.

Habitat indicators for SONEC shorebirds were evaluated using wetland trends in closed basin lakes. While seasonal and temporary wetland abundance increased substantially in these sites (Table S2, Figure S5), habitat impacts were characterized as moderate to acknowledge concerns about long-term ecosystem sustainability linked to accelerated patterns of lake drying shown by semi-permanent wetland loss (sensu Senner et al., 2018). In the Central Valley, semi-permanent, seasonal, and temporary wetland declines on duck clubs and wildlife refuges were indicators of significant shorebird migration and breeding (Apr-Sep) habitat impacts. Impacts to wintering shorebird (Nov-Mar) habitat in the Central Valley were moderate due to declining semi-permanent wetland abundance in combination with stable to increasing flooded agriculture. February was a significant outlier because of additional temporary wetland loss. Stable to increasing wetland trends in October showed only minor overlap with wintering shorebirds.

Moderate impacts were attributed to American bittern and white-faced ibis for most of their migration and wintering periods (Oct-Mar) in the Central Valley due to the loss of semi-permanent wetlands (Figure 9). Outliers included stable conditions in October and significant impacts in February that resulted from declines in semi-permanent and temporary wetlands. In SONEC, declining semi-permanent wetlands during breeding and summering periods (Apr-Sep) resulted in moderate habitat impacts five out of six months (Figure 9). Significant impacts occurred in July when declines occurred across all wetland types in addition to flooded agriculture. Breeding and summering impacts in the Central Valley were significant due to universal wetland declines from April to August. Significant impacts in September were due to reductions in temporary wetlands and flooded agriculture.

## 4.0 Discussion

Our analysis was the first we are aware of using a diverse suite of waterbird species as a framework for examining seasonal effects of wetland change within a flyway habitat network. Although linkages between wetlands and waterbirds were casual, results provide detailed insight into complex ecological trends and their relationship to interdependent life-cycle events.

Network habitats were provided by aggregating flooded agriculture and public-private wetland resources, including wildlife refuges. Declining wetland trends overlapping key breeding, migration, and wintering events were indicators of system-wide habitat declines, aligning in part with 33 waterbird species. This multi-species approach demonstrates the emergence of ecological risks through an improved understanding of wetland and waterbird interactions. Patterns of rapid wetland decline suggest that migratory networks in western North America may be approaching an ecological tipping point limiting their ability to support waterbird populations.

In SONEC and the Central Valley, pervasive loss of semi-permanent wetlands were indicators of functional decline that limited the availability of waterbird habitats. Losses resulted from shortened hydroperiods caused by excessive drying that forced the transition of semi-permanent to seasonal and temporary hydrologies—a process that in part offset concurrent seasonal and temporary wetland declines. Under this scenario, semi-permanent wetlands acted as a top-down index of ecosystem water balance decline due to their position at the top of the hydroperiod continuum. Similar patterns of functional decline have been observed in prairie and high-elevation wetland ecosystems that link accelerated drying to warming temperatures induced by climate change (McMenamin et al., 2008; Johnson et al., 2010; Lee et al., 2015).

Ecological effects that favor seasonal and temporary wetland availability were reinforced by flooded agriculture that mimicked shallow, intermittent surface water habitat in SONEC and the Central Valley. High proportional abundance and resilience of flooded agriculture worked in conjunction with top-down functional declines in semi-permanent systems as an additional buffer to seasonal and temporary wetland losses and were a major determinant of habitat availability. For example, in the Central Valley, favorable fall-winter habitat conditions were driven by flooded rice fields, which increased by 28% to 78% from November to January and were by far the largest contributor to waterbird habitat availability (sensu Fleskes et al., 2018). Likewise, reliable flood irrigation of grass hay from February to April has resulted in stable surface water conditions that currently account for 60% of available dabbling duck habitat during spring migration in SONEC (Donnelly et al., 2019).

Persistent summer loss of seasonal and temporary wetlands outside closed basin lakes was indicative of expanding top-down patterns of functional decline. Trends suggest that some functional groups have reached a point where increased evaporative demands during summer now outpace masking effects from the transformation of semi-permanents to seasonal and temporary hydroperiods. These patterns were most pronounced on public lands in SONEC (e.g., National Forest), where seasonal wetlands declined between 19% and 63% from May through August. Changes in water use priorities and/or policies may have also exacerbated declines on duck clubs and wildlife refuges that rely on artificial flooding to actively manage wetland conditions (Rosen et al., 2009). In SONEC, wetland availability on wildlife refuges has been impacted by the reallocation of limited water supplies in support of mandates to protect endangered fish species (Doremus and Tarlock, 2003). Additionally, the increased prevalence of mosquito-borne disease in the Central Valley has raised concerns over public safety (Githeko et al., 2000), leading to abatement measures that can significantly increase wetland management costs. Although the influence of mosquito control measures has not been quantified, they likely compound impacts of wetland declines because delayed flooding or intentional draining of wildlife refuges and duck clubs offers resource managers a low-cost solution to public health compliance (Berg et al., 2010).

### 4.1 Waterbird implications

Our results identified a clear concentration of impacts for waterbird species dependent on semi-permanent wetlands (Figure 10). Diving ducks, black terns, and grebes showed the greatest potential impact due to heavy use of semi-permanent wetlands, including littoral-limnetic systems occurring in closed-basin lakes, that support their primary habitat niche. Unlike other waterbirds evaluated, these species faced distinct challenges due to the ubiquitous nature of semi-permanent wetland loss that extended potential impacts across entire annual life cycles. Moreover, the effects of these impacts were amplified by a limited habitat base that omitted agriculturally supported habitats. Although agriculture has played an essential role in providing habitat that has offset historical wetland loss (Fasola and Ruiz, 1996; Elphick and Oring, 2003; Gauthier et al., 2005; Fox et al., 2017), it has contributed little to semi-permanent systems requiring some waterbird species to rely solely on wildlife refuges and remaining natural wetland resources to meet habitat needs.

**Figure 10.**
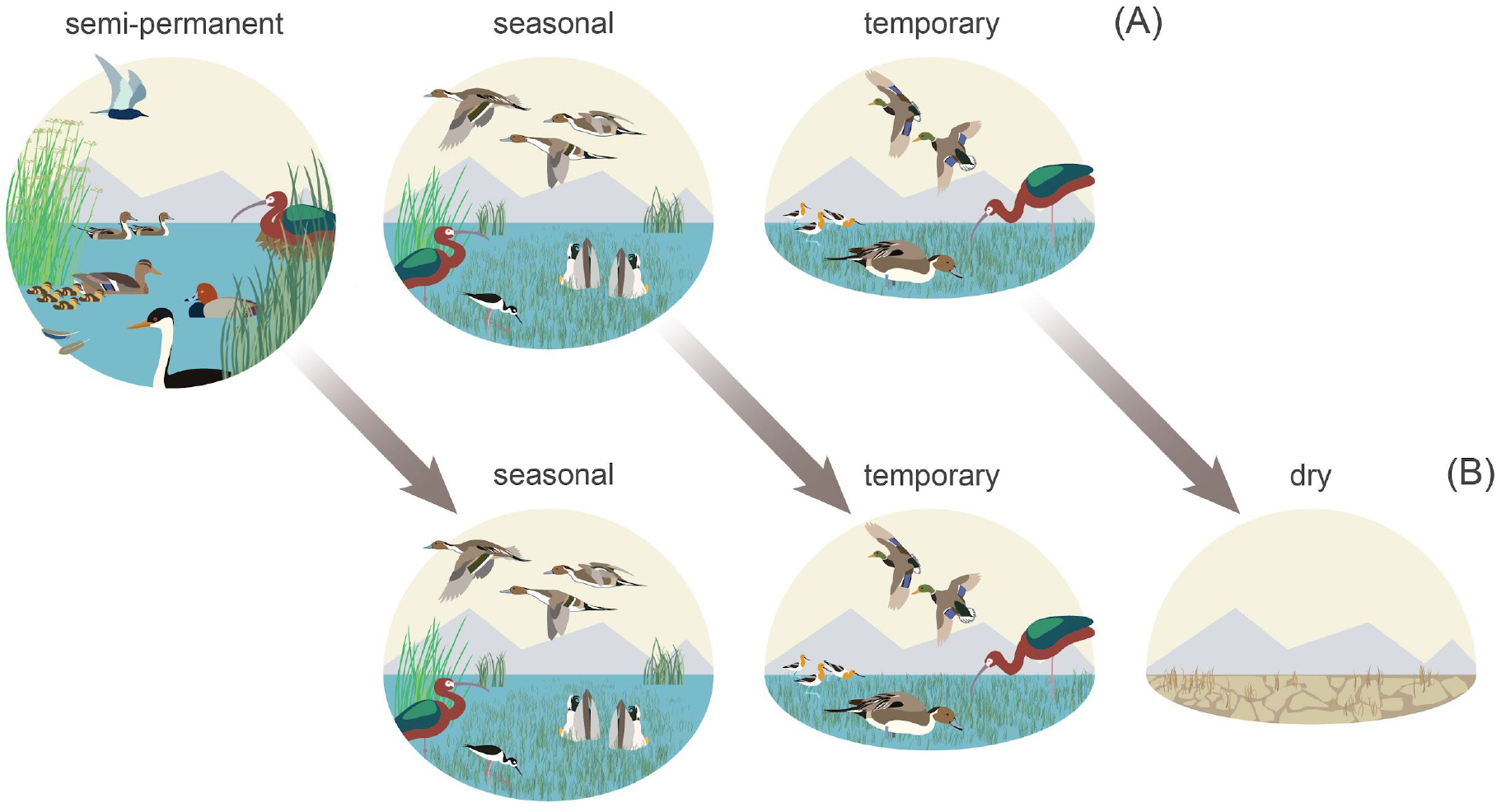
Functional wetland declines indicate disproportionate impacts to waterbird species heavily reliant on semi-permanent wetlands during all or portions of their annual life-cycle. Diving ducks (redhead), black terns, and grebes (western grebe) showed the greatest potential impact in addition to nesting white-faced ibis and molting and breeding waterfowl (A). Semi-permanent losses resulted from shortened hydroperiods caused by excessive drying that forced the transition of these habitats to seasonal and temporary hydologies—a process that offset concurrent seasonal and temporary wetland declines. Shorebirds (American avocets and black-necked stilts), migrating-wintering dabbling ducks (northern pintails and mallards), and white-faced ibis benefited from more persistent seasonal and temporary wetlands that were bolstered by stable agricultural habitats (B).

Wintering and migrating dabbling ducks represented one of our analysis’s least impacted habitat relationships (Figure 10). From October to April, birds benefited from relatively stable migration and wintering conditions in SONEC and the Central Valley. Conditions resulted from ecological trends, land-use, and management priorities on wildlife refuges and duck clubs that minimized impacts through a greater abundance of flooded agriculture (i.e., rice) and stable seasonal and temporary wetlands. Relationships were more complex for non-migratory dabbling ducks (i.e., cinnamon teal, gadwall, and mallard) that capitalized on reliable wintering conditions but were dependent on declining semi-permanent wetlands as breeding and molting habitat from April to September. Regionally declining cinnamon teal, gadwall, and mallard populations (Feldheim et al., 2018; USFWS 2020) and more persistent disease outbreaks may reflect impacts of degraded wetland conditions. In 2020, for example, ∼60,000 molting waterfowl were lost on a single wildlife refuge in SONEC due to botulism attributed to warming water temperatures and declining semi-permanent wetland abundance that concentrates birds in limited habitats (Sabalow, 2020).

Near-term effects of functional declines are less likely to impact species reliant on seasonal and temporary wetlands (Figure 10). While our results showed fewer impacts to these systems, their long-term sustainability remains uncertain. Loss of littoral-lacustrine wetland systems in SONEC closed-basin lakes, for example, has resulted in the exponential growth of seasonal and temporary wetlands that has increased habitat availability for some species. This is vividly illustrated at Goose Lake in SONEC, which now functions as one of the most extensive seasonal wetlands in the Pacific Flyway (Figure 11). However, rapid drying of littoral-lacustrine wetland systems in SONEC saline lakes (e.g., Abert and Summer) raises concerns over trophic collapse due to increased salinity associated with lower water volumes. Higher salinity can drastically reduce the diversity and biomass of benthic macroinvertebrates that serve as critical food resources for shorebirds and eared grebes. As water volumes continue to decrease, lakes can reach a point of infertility well before they dry entirely (Herbst, 2006; Moore, 2016; Senner et al., 2018). The transition of some declining freshwater lakes to saline states (sensu Thomas, 1995) may open habitat niches that offset losses in others. However, these lakes may also be vulnerable to collapse from salinity increases if lacustrine losses continue.

**Figure 11.**
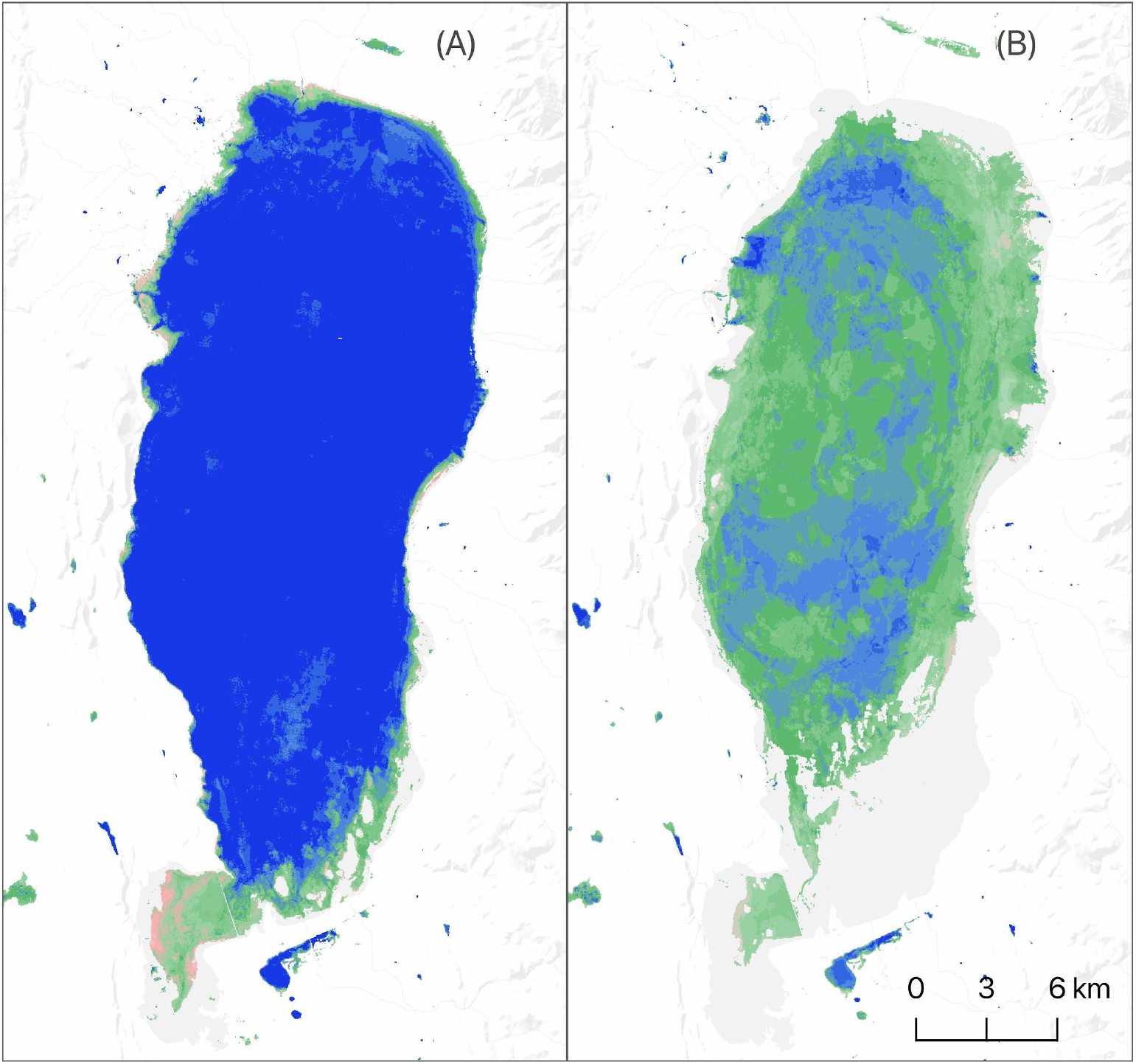
Goose Lake surface water and wetland hydroperiod extent [June] 1997 (A) and 2016 (B). Conditions representative of top-down functional transformation shown as drying littoral-limnetic systems in closed-basin lakes that lead to increased seasonal wetland abundance. Hydroperiods are defined by annual length of flooding: blue—semi-permanent (flooded > 8 months), green—seasonal (flooded > 2 and ≤ 8 months), and pink—temporary (flooded ≤ 2 months). Darker color shades indicate longer periods of inundation within hydroperiod classes.

Declining wetland trends on wildlife refuges and duck clubs from April to September were indicators of breeding shorebird impacts in the Central Valley. This region supports 24% and 17% of the U.S. breeding populations of American avocets and black-necked stilts, respectively (Shuford et al., 2007). Although most of these birds are known to breed in abundant flooded rice fields during spring (Shuford et al., 2007), conservation priorities identify the availability of wetlands on wildlife refuges and duck clubs as a vital factor sustaining habitat needs (USFWS 2020). However, current wetland trends suggest that it is unlikely wildlife refuges and duck clubs have the ability to provide breeding shorebird habitat due to management objectives that prioritize limited water supplies for wintering waterfowl. Alternative solutions include emerging conservation incentive programs that work with agricultural producers to flood fields on private lands as a stopgap measure to overcome shorebird habitat deficits (Reynolds et al., 2017).

### 4.2 Conservation needs

Impacts to waterbird migration networks identified in this study represent the early effects of climate change. A posthoc analysis of drought indices for both SONEC and the Central Valley (*see* Supplemental Materials - Recent Climate) identified intensifying patterns of drought over the study period. Changes were most pronounced in SONEC, where drought has become the regional norm since 2005 (Figure S11). Our findings suggest that drought effects are ubiquitous and can impact wetland function regardless of underlying hydrologic mechanisms (e.g., managed or natural). The Central Valley, for example, relies on reservoir storage capacity 22 times greater than SONEC to attenuate drought by storing snow-melt runoff to provide water for agriculture and artificially managed wetlands (Table S11). Although these systems were developed to ensure reliable water supplies, higher frequency and more severe drought events (Diffenbaugh et al., 2015; Swain, 2021) have triggered measures curtailing water deliveries to wildlife refuges (Rosen et al., 2009) that have mirrored more direct ecological effects of wetland loss within SONEC (Donnelly et al., 2020).

While the stability of agriculturally supported wetlands implies potential climate resilience, they are more vulnerable to indirect economic pressures related to increasing water scarcity that can significantly reduce wildlife benefits (Mann and Gleick, 2015). Potential impacts are greatest in the Central Valley, where many waterbird species have become dependent on flooded agriculture (primarily flooded rice) that makeup ∼75% of the region’s habitat annually. Winter flooding of rice fields to remove post-harvest stubble was initially triggered by the Federal Clean Air Act and subsequent California state legislation in 1991 that mitigated historic burning practices. Abundant water resources for winter decomposition of rice stubble (a boon for wetland habitats) offered an economically viable solution to burning. Our results showed producer adoption of this technique increased winter availability of agricultural habitats by as much as 78%, making it an indispensable component of the migratory network that has translated to higher waterbird survival and forage capacity (Fleskes et al., 2007, 2016; Strum et al., 2013). While we found minimal evidence of declining rice cultivation overall (<4%), new economic incentives for rice straw used in fiber-board manufacturing are providing producers alternatives to winter flooding as the reliability of irrigation water declines (Gibson, 2019).

While our analysis did not measure surface and groundwater interactions directly, groundwater sustainability is crucial to maintaining surface water hydrology in most wetland ecosystems, particularly in arid and semi-arid regions in western North America (sensu Jolly et al., 2008). Recent work from Thomas et al. (2017) and Wang et al. (2016) identify clear linkages between intensifying meteorological drought and reduced groundwater storage. Moreover, Kibler et al. (2021) found that dieback of riparian vegetation (dependent on shallow alluvial aquifers) was a direct result of depleted groundwater during the 2012-19 California drought. Compounding declines are shifts in agricultural water consumption in SONEC and the Central Valley that increasingly rely on groundwater extraction as a primary irrigation source to offset surface water declines of ∼30% over the past decade (Medellín-Azuara et al., 2015). Climate scenario planning to maintain agricultural production in the Central Valley has identified conversion to more profitable and water-saving crops as a viable solution that supports economic viability and recovers groundwater depletions to alleviate drought (Li et al., 2018). Indirect benefits of such actions may improve climate resilience in some wetland systems. Still, they may also result in a net loss of agricultural habitat by reducing water-intensive crops like rice that currently support large waterbird populations.

There was little indication that changing agricultural practices resulted in waterbird habitat loss in SONEC. Similar regions in the western United States, however, are under increasing pressure from climate-driven initiatives to adopt more efficient irrigation technology (e.g., center pivot sprinkler irrigation) and rotational fallowing that would transfer water savings to municipal use (Thorvaldson and Pritchett, 2006; Welsh and Endter-Wada, 2017). While these efforts seek viable solutions to climate change and urban water demands, they often disregard ecosystem services associated with flooded agriculture. For example, the common practice of flood irrigating grass hay (occurring predominantly in riparian floodplains, Donnelly et al., 2020) mimics once natural hydrologic processes. Still, it is frequently deemed an inefficient use of water (Richter et al., 2017). Instead, these practices have been shown to promote climate resiliency through groundwater recharge that generates late summer return flows in adjacent streams, benefiting waterbirds, fisheries, and riparian habitats (Blevins et al., 2016). Future protections of agriculturally supported wetlands in SONEC will likely require a better understanding of ecological tradeoffs associated with water reallocation as the need for climate change adaptations rise.

Climate forcing will likely continue to reshape SONEC and Central Valley wetland ecosystems. Recent projections from Snyder et al. (2019) show that by 2020-2050 regional temperatures will be ∼1°C to ∼3°C above the historical baseline of 1980-2010. More importantly, Cook et al. (2015) showed that rising temperatures driving increased evapotranspiration would lead to ‘unprecedented’ drought throughout the region. Our posthoc analyses of downscaled future climate data for SONEC and the Central Valley show a more intense and continuous drought (*see* Supplemental Materials - Future Climate). Projected changes are likely to force tradeoffs in water use priorities that could intensify ecological risks already identified in our analysis. Under these scenarios, it will become increasingly important to consider adaptations that preserve ecological and anthropogenic (e.g., flooded agriculture) mechanisms supporting wetland resilience. Emerging solutions include increased recognition of ecosystem services provided through beneficial agricultural practices by giving producers economic incentives to maintain flood irrigation. Recent efforts include a program in the Central Valley that uses winter-flooded rice fields (supporting waterbirds) to rear endangered chinook salmon *(Oncorhynchus tshawytscha)* smolt to increase fish survival (Holmes et al., 2021). In other regions of the western U.S., groups are exploring conservation exchange programs to establish a market for private investment in ecosystem services that will pay ranchers for maintaining flood irrigation practices in grass hay meadows that are mutually beneficial to wildlife and riparian sustainability (Duke et al., 2011; Blevins et al., 2016).

Conservation strategies that preserve climate resiliency must also consider adaptive measures needed to maintain overall flyway function. Intensifying water scarcity during future droughts could change the roles of SONEC and the Central Valley as waterbirds seek more productive landscapes to support stopover and wintering needs. Donnelly et al. (2020) identified nonlinear patterns of wetland drying in North American waterbird flyways that showed significant wetland impacts to snowmelt-driven systems in the western U.S., while monsoon-driven wetlands that overlap wintering waterbird distributions in Mexico remained stable or expanded over time. Migratory waterbirds are well adapted to take advantage of shifting continental conditions and have shown an ability to alter habitat use within flyways as climate change restructures resource availability (Lehikoinen et al., 2013; Pavón-Jordán et al., 2015). Under these scenarios, resource managers must be willing to proactively prioritize and adapt management strategies that reflect an evolution in waterbird habitat needs, including redirection of conservation investments to more resilient regions of the flyway that are likely to support future waterbird populations.

Balancing specific social, ecological, and economic factors will be necessary to accurately identify trade-offs affecting wetlands and the resiliency of waterbird migration networks. This study highlights that waterbird impacts are manifested through complex interactions between interdependent landscapes that experience independent habitat risks. Increased pressure on waterbird migration networks will require increased coordination between important waterbird breeding, wintering, and stopover regions to proactively identify and address emerging risks impacting populations as changes to climate and land use accelerate. To inform wetland and waterbird conservation, we make our data available through an interactive web-based application allowing natural resource managers direct access to long-term wetland trends used in our analysis (insert link). We encourage using our findings to inform solutions to wetland loss through collaborative and proactive decision-making among local and regional stakeholders throughout waterbird flyways of western North America.

## Data Availability Statement

The original data presented in the study are publicly available and can be found here: (insert link)

## Author Contributions

JPD conceived and designed the study. JPD and JM conducted the wetland and waterbird analysis. JPD and JM wrote the manuscript. MC and SC contributed to the manuscript. All authors contributed to the article and approved the submitted version.

## Acknowledgments

We thank John Vradenburg, senior biologist - Klamath National Wildlife Refuge Complex, and Michael D’Errico senior biologist - Sacramento Refuge complex for their insight and waterfowl and shorebird survey data that supported this analysis. We also thank the U.S. Fish and Wildlife Service for funding that made this work possible. Views in this manuscript from U. S. Fish and Wildlife Service authors are their own and do not necessarily represent the agency’s views. Any use of trade, firm, or product names is for descriptive purposes only and does not imply endorsement by the U.S. Government.

## Supplemental Material

See document

