## supplemental material for "Functional wetland loss drives emerging risks to waterbird migration networks"

**Supplemental Materials - Methods**

**Methods Section 1 - remote sensing methods for wetland hydrology trends**

Following the methods that Donnelly et al. (2021) outlined, wetland and agricultural surface water conditions were measured monthly as a five-year running means using constrained spectral mixture analysis (SMA; Adams and Gillespie 2006). This approach allowed proportional estimations of water contained within a continuous 30×30 m pixel grid (Halabisky *et al.* 2016; Jin et al. 2017) and provided an accurate account of flooding when detectability was reduced due to interspersion of emergent vegetation, shallow, or turbid water (DeVries *et al.* 2017), characteristics common to seasonal wetlands in semi-arid regions (Jolly *et al.* 2008). Because these conditions can partially mask areas covered with water (Donnelly *et al.* 2019), we considered pixels fully inundated when water was present. Pixels containing <15% surface water were omitted from summaries to minimize the overestimation of surface water area.

Satellite data used for SMA were formatted by binning individual Landsat scenes by month and averaging results into twelve composite images for each five-year mean. Results provided 444 unique monthly measures of wetland-agriculture surface water for the SONEC and Central Valley regions. Areas containing cloud, cloud shadow, snow, and ice were masked using the Landsat CFMask band (Foga *et al.* 2017). All unmasked pixels in Landsat 30 m visible, near-infrared, and short wave infrared bands were incorporated into SMA except for Landsat 8 coastal aerosol band. Surface water was not measured in 2012 due to poor quality satellite imagery.

Training data for SMA were extracted from satellite imagery as spectral end members unique to individual images classified. Training site locations represented homogeneous land cover types mapped as water, wetland vegetation, upland, and alkali soil. Spectral end members for water were collected using image masks generated from 99th percentile normalized difference water index values (McFeeters 1996), coincident with large deepwater lakes within both regions. A similar masking approach was applied to collect wetland vegetation end members using normalized difference vegetation indices (Box *et al.* 1989). Sampling was constrained to sites coincident with flooded wetlands and representative of associated plant phenology. Spectral mixture analysis requires minimal training data (Adams and Gillespie 2006) that allows upland and alkali soil end members to be generated from a small number of static plots within the regions (n = 4; 0.5-1 $km^2$). Upland plots were associated with homogenous shrublands characterized by low vegetative productivity and high soil exposure. Alkali soil plots were coincident with dry lake basins in surface mineral deposits. Plot locations were identified using high resolution (< 0.5 m) multispectral satellite imagery or field survey. All image processing and raster‑based analyses were conducted using Google Earth Engine cloud‑based geospatial processing platform (Gorelick *et al.* 2017).

**Supplemental Section 2 - change detection methods for wetland loss**

Change detection analysis was used to designate wetland or flooded agricultural declines as functional or physical loss to discern underlying drivers of change. Functional losses were attributed to areas of diminishing surface water (i.e. drying) associated with shifts in ecological water balance or water management in the absence of physical alterations. Land conversion (e.g., urban expansion or shifting agricultural practices) resulting in surface water declines were identified as physical loss. Areas of change were delineated by differencing mean monthly (Jan-Dec) surface water conditions between T1 (1984-1991) and T2 (2013-20). Using a GIS, change areas were visually inspected through on-screen photo-interpretation of high resolution (≤ 1 m) multispectral satellite imagery (acquired 2018 or later) to identify areas of physical loss. Surface water conditions for T1 and T2 were derived using remote-sensing methods outlined in Supplemental Section 1. All image processing and raster‑based analyses were conducted using Google Earth Engine cloud‑based geospatial processing platform (Gorelick *et al.* 2017). GIS analyses were performed using QGIS (QGIS Development Team 2020).

**Supplemental Section 3 - eBird-traditional survey comparison**

To compare temporal abundance distributions derived from the eBird Basic Dataset (EBD) and traditional survey methods (i.e., aerial survey and systematic ground counts), waterbird counts were binned bi-weekly and summed across years. Results were then grouped by region, species, and survey type and scaled to relative values. Boxplots and non-parametric Wilcoxon tests were used to display and compare data graphically. All available EBD observations collected from 1984 to 2020 in the SONEC and Central Valley regions were used in our evaluation. SONEC bi-weekly aerial waterfowl surveys conducted in the Klamath Basin from 1984-2016 were used for EBD evaluation. Because surveys were flown during spring (Jan-May) and fall (Sep-Dec), distributions were compared for each period using four migrating dabbling duck species (Fig. S1-2). Although aerial survey efforts were conducted for a subset of SONEC, results were considered representative of regional waterfowl use patterns (Donnelly *et al.* 2019).

Bi-weekly ground surveys on the Sacramento National Wildlife Refuge Complex (hereafter 'refuge complex') were used in the Central Valley for EBD evaluation. Ground surveys were collected from 2011 to 2017 across six independent refuge units representing the northern half of the Central Valley study area. For comparison, we selected five wintering waterfowl (Figs. S3) and three fall migrating shorebird species (Fig. S4) based on their use of habitats associated with the refuge complex. SONEC and Central Valley comparisons showed no significant differences in temporal abundance patterns. Outcomes support previous results from Callaghan and Gawlik (2015) and Walker and Taylor (2017) that showed EBD observations and traditional survey efforts equivalent when applied at broad scales.

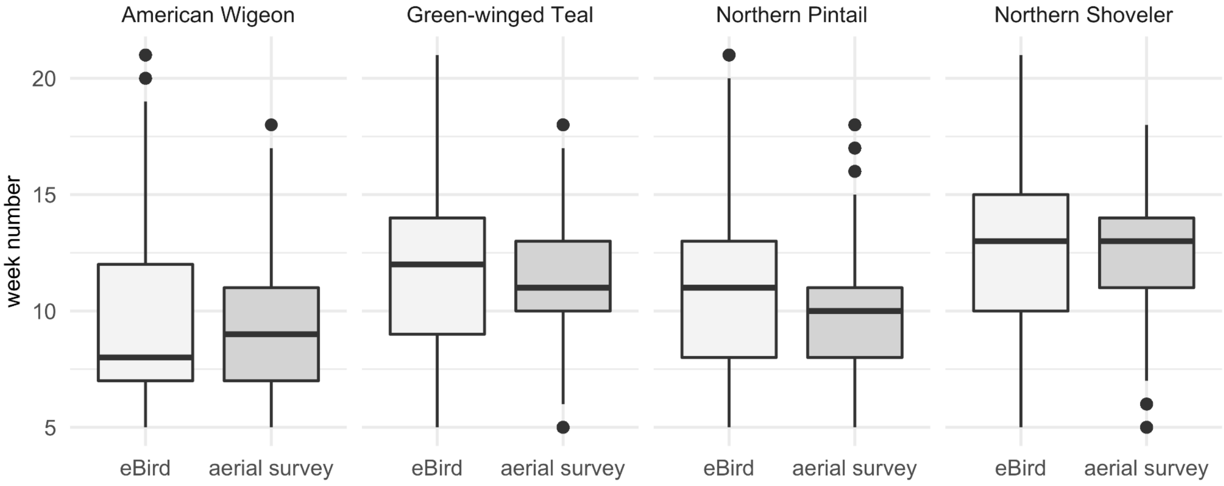


**Fig. S1.** Temporal distribution of dabbling duck abundance derived from eBird Basic Dataset and aerial surveys collected during spring migration (Feb-May) in SONEC. Distributions representative of all available eBird (1984-2020) and aerial survey counts (1984 to 2016). Nonparametric Wilcoxon tests results by species: American Wigeon p-value 0.258, Green-winged Teal p-value 0.776, Northern Pintail p-value 0.315, Northern Shoveler p-value 0.972.

temporal distribution of dabbling duck abundance derived from

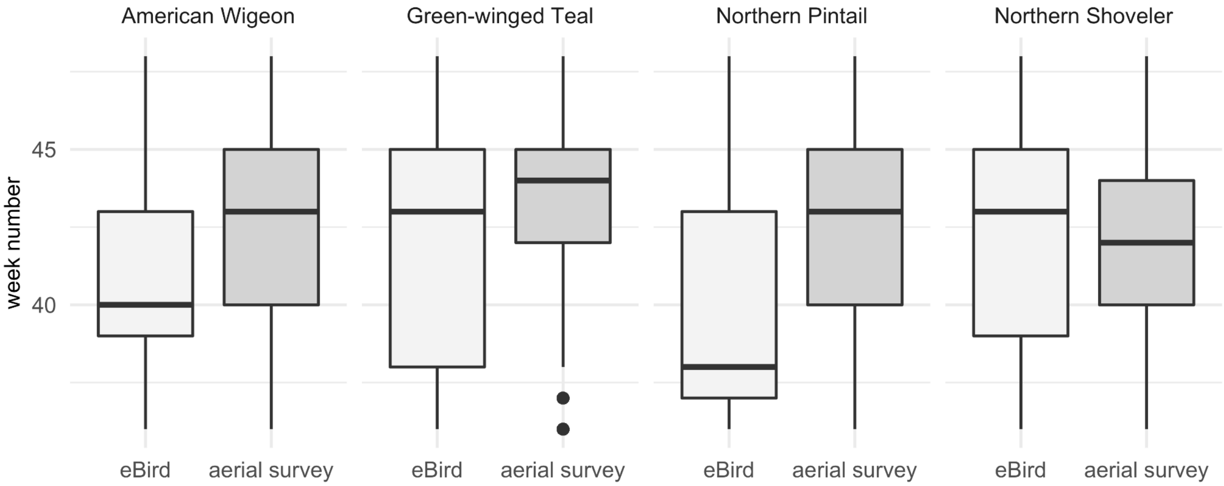


**Fig. S2**.Temporal distribution of dabbling duck abundance derived from eBird Basic Dataset and aerial surveys collected during spring migration (Sep-Dec) in SONEC. Distributions representative of all available eBird (1984-2020) and aerial survey counts (1984 to 2016). Nonparametric Wilcoxon tests results by species: American Wigeon p-value 0.258, Green-winged Teal p-value 0.776, Northern Pintail p-value 0.315, Northern Shoveler p-value 0.972.

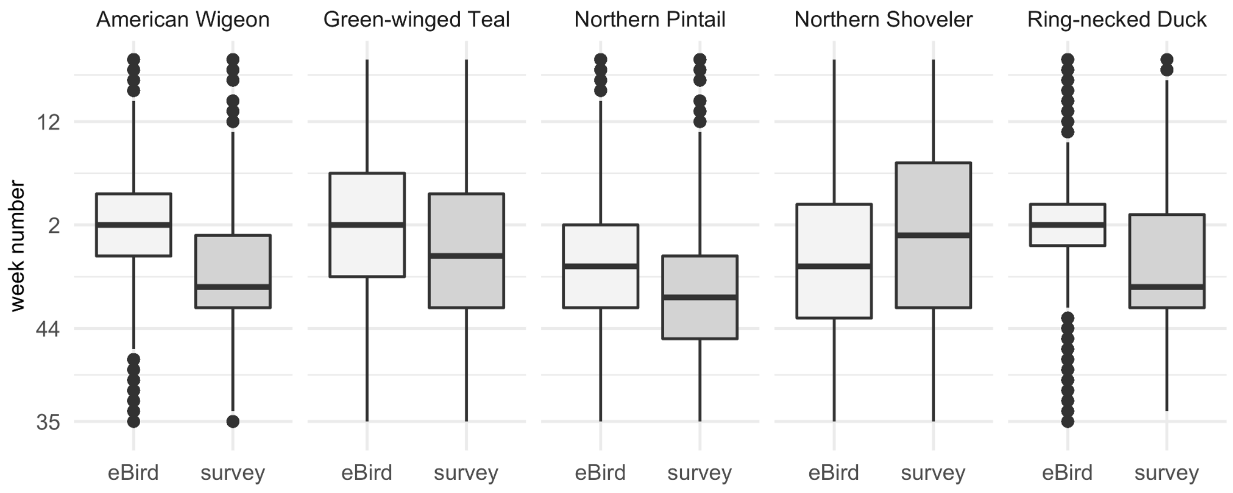


**Fig. S3**. Temporal distribution of dabbling duck abundance derived from eBird Basic Dataset and ground surveys collected during the wintering period (Oct-Mar) in the Central Valley. Distributions representative of all available eBird (1984-2020) and ground survey counts (2011 to 2017). Nonparametric Wilcoxon tests results by species: American Wigeon p-value 0.480, Green-winged Teal p-value 0.893, Northern Pintail p-value 0.757, Northern Shoveler p-value 0.941, Ring-necked Duck p-value 0.628.

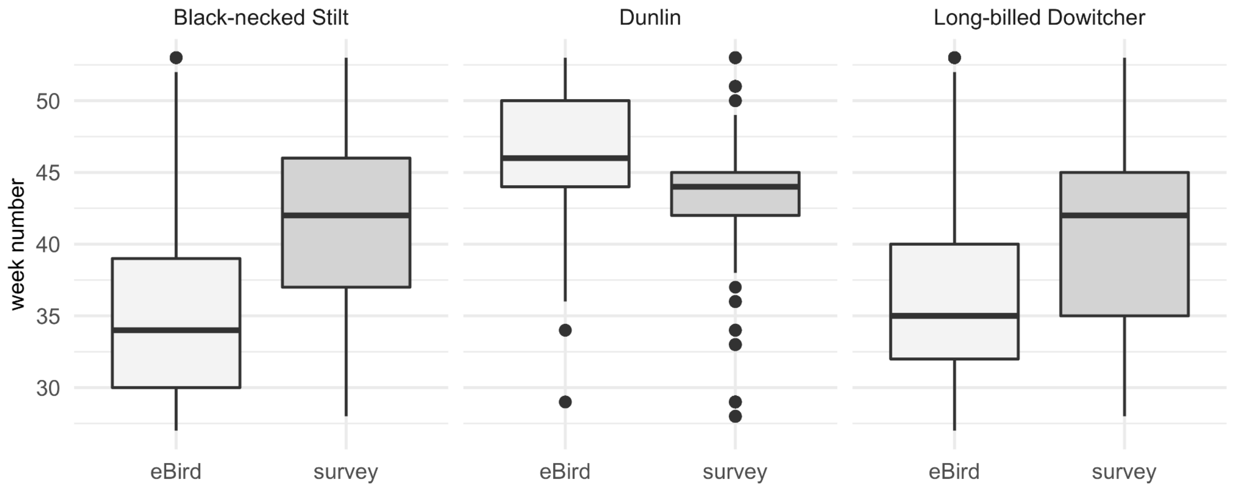


**Fig. S4.** Temporal distribution of shorebird abundance derived from eBird Basic Dataset and ground surveys collected from August to December in the Central Valley (see Fig. 1). Distributions representative of all available eBird (1984-2020) and ground survey counts (2011 to 2017). Nonparametric Wilcoxon tests results by species: Black-necked Stilt p-value 0.968, Dunlin p-value 0.072, Long-billed Dowitcher p-value 0.698.

**Supplementary Materials -- Results**

Table S1: SONEC all wetlands - P1 (1988-2004) and P2 (2005-20) median monthly surface water change. Areas (ha) include wetlands associated with closed basin lakes, public-private lands, and wildlife refuges.

| Hydroperiod | Month | P1 (1988-2004) | P2 (2004-2020) | Difference | % Difference | Wilcox *p* |
|---|---|---|---|---|---|---|
| semi-permanent | Jan | 183,964 | 89,818 | -94,147 | -51% | 0.000 |
| | Feb | 189,984 | 122,833 | -67,151 | -35% | 0.002 |
| | Mar | 213,765 | 156,377 | -57,388 | -27% | 0.000 |
| | Apr | 220,250 | 160,551 | -59,699 | -27% | 0.000 |
| | May | 246,637 | 157,925 | -88,712 | -36% | 0.000 |
| | Jun | 242,978 | 162,556 | -80,422 | -33% | 0.000 |
| | Jul | 243,206 | 152,021 | -91,185 | -37% | 0.000 |
| | Aug | 229,178 | 144,744 | -84,434 | -37% | 0.000 |
| | Sep | 216,979 | 140,497 | -76,482 | -35% | 0.000 |
| | Oct | 216,636 | 117,410 | -99,226 | -46% | 0.000 |
| | Nov | 194,011 | 113,630 | -80,381 | -41% | 0.000 |
| | Dec | 187,751 | 110,171 | -77,580 | -41% | 0.000 |
| seasonal | Jan | 22,687 | 18,139 | -4,547 | -20% | 0.102 |
| | Feb | 36,228 | 39,209 | 2,980 | 8% | 0.204 |
| | Mar | 41,984 | 50,821 | 8,836 | 21% | 0.004 |
| | Apr | 39,804 | 52,155 | 12,351 | 31% | 0.001 |
| | May | 37,862 | 51,196 | 13,334 | 35% | 0.002 |
| | Jun | 33,661 | 45,951 | 12,290 | 37% | 0.001 |
| | Jul | 22,837 | 16,408 | -6,429 | -28% | 0.023 |
| | Aug | 5,562 | 3,845 | -1,717 | -31% | 0.191 |
| | Sep | 2,849 | 2,993 | 144 | 5% | 0.998 |
| | Oct | 4,651 | 7,428 | 2,778 | 60% | 0.001 |
| | Nov | 10,793 | 12,104 | 1,312 | 12% | 0.049 |
| | Dec | 18,119 | 18,716 | 598 | 3% | 0.873 |
| temporary | Jan | 27,519 | 40,603 | 13,084 | 48% | 0.136 |
| | Feb | 56,432 | 55,041 | -1,390 | -3% | 0.606 |
| | Mar | 37,641 | 39,312 | 1,671 | 4% | 0.709 |
| | Apr | 23,915 | 32,674 | 8,759 | 37% | 0.191 |
| | May | 19,081 | 31,081 | 12,000 | 63% | 0.000 |
| | Jun | 13,315 | 15,191 | 1,876 | 14% | 0.204 |
| | Jul | 3,986 | 2,054 | -1,932 | -49% | 0.008 |
| | Aug | 388 | 605 | 217 | 56% | 0.127 |
| | Sep | 385 | 509 | 123 | 32% | 0.045 |
| | Oct | 987 | 2,028 | 1,041 | 106% | 0.000 |
| | Nov | 9,116 | 9,372 | 256 | 3% | 0.465 |
| | Dec | 24,954 | 23,567 | -1,387 | -6% | 0.444 |

Table S2: SONEC closed basin lakes - P1 (1988-2004) and P2 (2005-20) median monthly surface water change. Areas (ha) inclusive of all littoral-lacustrine wetland systems.

| Hydroperiod | Month | P1 (1988-2004) | P2 (2004-2020) | Difference | % Difference | Wilcox *p* |
|---|---|---|---|---|---|---|
| | Jan | 132,014 | 62,045 | -69,969 | -53% | 0.000 |
| | Feb | 147,516 | 88,625 | -58,891 | -40% | 0.001 |
| | Mar | 157,163 | 113,450 | -43,712 | -28% | 0.000 |
| | Apr | 167,473 | 115,991 | -51,481 | -31% | 0.000 |
| | May | 183,108 | 120,152 | -62,956 | -34% | 0.000 |
| semi-permanent | Jun | 182,999 | 122,005 | -60,994 | -33% | 0.000 |
| | Jul | 182,902 | 117,643 | -65,259 | -36% | 0.000 |
| | Aug | 178,617 | 114,286 | -64,331 | -36% | 0.000 |
| | Sep | 169,799 | 112,115 | -57,685 | -34% | 0.000 |
| | Oct | 167,106 | 88,037 | -79,069 | -47% | 0.000 |
| | Nov | 151,571 | 84,091 | -67,481 | -45% | 0.000 |
| | Dec | 145,878 | 85,281 | -60,597 | -42% | 0.000 |
| | Jan | 4,298 | 6,733 | 2,434 | 57% | 0.326 |
| | Feb | 8,231 | 11,888 | 3,657 | 44% | 0.025 |
| | Mar | 6,499 | 17,371 | 10,872 | 167% | 0.005 |
| | Apr | 9,385 | 23,608 | 14,223 | 152% | 0.000 |
| | May | 11,409 | 24,169 | 12,760 | 112% | 0.000 |
| seasonal | Jun | 10,476 | 22,618 | 12,142 | 116% | 0.000 |
| | Jul | 8,661 | 8,502 | -159 | -2% | 0.845 |
| | Aug | 2,099 | 2,240 | 141 | 7% | 0.763 |
| | Sep | 864 | 1,621 | 757 | 88% | 0.245 |
| | Oct | 1,120 | 2,574 | 1,454 | 130% | 0.001 |
| | Nov | 2,322 | 5,976 | 3,654 | 157% | 0.000 |
| | Dec | 4,569 | 7,818 | 3,250 | 71% | 0.058 |
| | Jan | 4,798 | 8,324 | 3,526 | 74% | 0.025 |
| | Feb | 7,649 | 12,314 | 4,664 | 61% | 0.045 |
| | Mar | 2,675 | 7,363 | 4,688 | 175% | 0.023 |
| | Apr | 3,647 | 12,830 | 9,183 | 252% | 0.001 |
| | May | 3,699 | 13,504 | 9,806 | 265% | 0.000 |
| temporary | Jun | 2,041 | 7,504 | 5,463 | 268% | 0.001 |
| | Jul | 1,122 | 498 | -624 | -56% | 0.245 |
| | Aug | 87 | 264 | 177 | 203% | 0.017 |
| | Sep | 93 | 205 | 111 | 119% | 0.003 |
| | Oct | 159 | 563 | 404 | 254% | 0.000 |
| | Nov | 855 | 1,944 | 1,089 | 127% | 0.009 |
| | Dec | 4,420 | 4,793 | 373 | 8% | 0.736 |

Table S3: SONEC wildlife refuges - P1 (1988-2004) and P2 (2005-20) median monthly surface water change. Areas (ha) exclusive to state and federally managed wildlife refuges.

| Hydroperiod | Month | P1 (1988-2004) | P2 (2004-2020) | Difference | % Difference | Wilcox *p* |
|---|---|---|---|---|---|---|
| semi-permanent | Jan | 12,870 | 8,714 | -4,156 | -32% | 0.000 |
| | Feb | 15,623 | 11,399 | -4,224 | -27% | 0.000 |
| | Mar | 15,566 | 12,578 | -2,989 | -19% | 0.000 |
| | Apr | 14,658 | 12,449 | -2,209 | -15% | 0.001 |
| | May | 15,107 | 11,128 | -3,979 | -26% | 0.000 |
| | Jun | 14,756 | 11,296 | -3,460 | -23% | 0.000 |
| | Jul | 13,063 | 9,765 | -3,298 | -25% | 0.000 |
| | Aug | 11,691 | 8,262 | -3,429 | -29% | 0.000 |
| | Sep | 11,348 | 7,803 | -3,544 | -31% | 0.000 |
| | Oct | 12,695 | 10,401 | -2,295 | -18% | 0.000 |
| | Nov | 13,898 | 10,884 | -3,015 | -22% | 0.000 |
| | Dec | 12,810 | 9,613 | -3,197 | -25% | 0.000 |
| seasonal | Jan | 6,466 | 4,054 | -2,412 | -37% | 0.031 |
| | Feb | 11,319 | 7,059 | -4,260 | -38% | 0.204 |
| | Mar | 11,844 | 13,278 | 1,434 | 12% | 0.309 |
| | Apr | 9,202 | 10,816 | 1,614 | 18% | 0.127 |
| | May | 5,709 | 7,595 | 1,886 | 33% | 0.034 |
| | Jun | 3,962 | 6,613 | 2,651 | 67% | 0.015 |
| | Jul | 1,836 | 1,887 | 51 | 3% | 0.873 |
| | Aug | 360 | 304 | -56 | -16% | 0.533 |
| | Sep | 378 | 335 | -42 | -11% | 0.231 |
| | Oct | 1,059 | 1,179 | 119 | 11% | 0.292 |
| | Nov | 3,093 | 2,243 | -850 | -27% | 0.017 |
| | Dec | 4,967 | 3,797 | -1,170 | -24% | 0.023 |
| temporary | Jan | 6,364 | 4,771 | -1,593 | -25% | 0.790 |
| | Feb | 13,003 | 8,342 | -4,660 | -36% | 0.005 |
| | Mar | 10,580 | 9,657 | -923 | -9% | 0.292 |
| | Apr | 5,986 | 4,498 | -1,488 | -25% | 0.488 |
| | May | 2,393 | 2,490 | 98 | 4% | 0.709 |
| | Jun | 1,186 | 1,997 | 812 | 68% | 0.025 |
| | Jul | 574 | 410 | -164 | -29% | 0.191 |
| | Aug | 55 | 93 | 38 | 68% | 0.008 |
| | Sep | 94 | 112 | 18 | 20% | 0.557 |
| | Oct | 280 | 353 | 72 | 26% | 0.402 |
| | Nov | 1,362 | 1,142 | -220 | -16% | 0.276 |
| | Dec | 5,264 | 3,740 | -1,524 | -29% | 0.146 |

Table S4: SONEC public wetlands - P1 (1988-2004) and P2 (2005-20) median monthly surface water change. Areas (ha) encompass un-managed or natural wetlands on public lands administered by, but not limited to the U.S. Forest Service and Bureau of Land Management.

| Hydroperiod | Month | P1 (1988-2004) | P2 (2004-2020) | Difference | % Difference | Wilcox *p* |
|---|---|---|---|---|---|---|
| | Jan | 13,525 | 5,361 | -8,164 | -60% | 0.008 |
| | Feb | 12,800 | 10,234 | -2,566 | -20% | 0.041 |
| | Mar | 18,875 | 11,332 | -7,543 | -40% | 0.000 |
| | Apr | 16,487 | 11,763 | -4,724 | -29% | 0.002 |
| | May | 18,244 | 11,030 | -7,214 | -40% | 0.000 |
| semi-permanent | Jun | 17,704 | 11,381 | -6,323 | -36% | 0.000 |
| | Jul | 17,971 | 10,184 | -7,787 | -43% | 0.000 |
| | Aug | 16,598 | 9,537 | -7,061 | -43% | 0.001 |
| | Sep | 13,358 | 9,085 | -4,273 | -32% | 0.000 |
| | Oct | 13,338 | 9,078 | -4,260 | -32% | 0.003 |
| | Nov | 13,564 | 9,434 | -4,131 | -30% | 0.000 |
| | Dec | 12,245 | 6,278 | -5,967 | -49% | 0.004 |
| | Jan | 4,785 | 2,501 | -2,284 | -48% | 0.034 |
| | Feb | 7,265 | 9,342 | 2,077 | 29% | 0.023 |
| | Mar | 9,660 | 10,191 | 531 | 6% | 0.790 |
| | Apr | 9,884 | 10,491 | 607 | 6% | 0.873 |
| | May | 11,063 | 8,999 | -2,064 | -19% | 0.074 |
| seasonal | Jun | 9,880 | 8,284 | -1,596 | -16% | 0.034 |
| | Jul | 7,611 | 2,811 | -4,800 | -63% | 0.000 |
| | Aug | 1,591 | 846 | -745 | -47% | 0.002 |
| | Sep | 783 | 568 | -215 | -27% | 0.231 |
| | Oct | 767 | 1,149 | 382 | 50% | 0.037 |
| | Nov | 2,161 | 2,612 | 452 | 21% | 0.231 |
| | Dec | 3,267 | 3,230 | -37 | -1% | 0.817 |
| | Jan | 5,234 | 7,199 | 1,965 | 38% | 0.326 |
| | Feb | 12,039 | 14,409 | 2,370 | 20% | 0.260 |
| | Mar | 8,669 | 7,578 | -1,091 | -13% | 0.292 |
| | Apr | 5,497 | 6,774 | 1,277 | 23% | 0.292 |
| | May | 4,987 | 5,849 | 863 | 17% | 0.709 |
| temporary | Jun | 3,961 | 3,353 | -608 | -15% | 0.309 |
| | Jul | 864 | 310 | -554 | -64% | 0.000 |
| | Aug | 68 | 101 | 32 | 47% | 0.631 |
| | Sep | 63 | 58 | -4 | -7% | 0.986 |
| | Oct | 100 | 336 | 236 | 237% | 0.000 |
| | Nov | 1,791 | 2,863 | 1,073 | 60% | 0.010 |
| | Dec | 5,398 | 6,972 | 1,574 | 29% | 0.736 |

Table S5: SONEC private wetlands - P1 (1988-2004) and P2 (2005-20) median monthly surface water change. Areas (ha) exclusive to private un-managed or natural wetlands.

| Hydroperiod | Month | P1 (1988-2004) | P2 (2004-2020) | Difference | % Difference | Wilcox *p* |
|---|---|---|---|---|---|---|
| semi-permanent | Jan | 8,322 | 6,285 | -2,037 | -24% | 0.110 |
| | Feb | 9,699 | 9,029 | -670 | -7% | 0.245 |
| | Mar | 12,909 | 11,154 | -1,755 | -14% | 0.045 |
| | Apr | 12,761 | 11,157 | -1,605 | -13% | 0.157 |
| | May | 12,691 | 9,564 | -3,127 | -25% | 0.045 |
| | Jun | 12,705 | 10,767 | -1,937 | -15% | 0.058 |
| | Jul | 12,502 | 9,748 | -2,754 | -22% | 0.015 |
| | Aug | 11,351 | 7,978 | -3,373 | -30% | 0.037 |
| | Sep | 9,676 | 7,006 | -2,670 | -28% | 0.058 |
| | Oct | 10,644 | 6,959 | -3,684 | -35% | 0.045 |
| | Nov | 10,236 | 7,343 | -2,893 | -28% | 0.004 |
| | Dec | 9,232 | 5,210 | -4,022 | -44% | 0.008 |
| seasonal | Jan | 5,046 | 3,315 | -1,731 | -34% | 0.510 |
| | Feb | 9,102 | 8,856 | -246 | -3% | 0.657 |
| | Mar | 9,819 | 11,080 | 1,261 | 13% | 0.136 |
| | Apr | 8,504 | 9,984 | 1,480 | 17% | 0.276 |
| | May | 8,726 | 8,768 | 42 | 1% | 0.901 |
| | Jun | 7,699 | 8,311 | 612 | 8% | 0.488 |
| | Jul | 4,862 | 3,316 | -1,546 | -32% | 0.000 |
| | Aug | 1,279 | 672 | -607 | -48% | 0.009 |
| | Sep | 588 | 428 | -159 | -27% | 0.292 |
| | Oct | 687 | 891 | 204 | 30% | 0.382 |
| | Nov | 2,527 | 1,889 | -638 | -25% | 0.087 |
| | Dec | 3,154 | 2,507 | -647 | -21% | 0.709 |
| temporary | Jan | 7,144 | 7,119 | -25 | 0% | 0.606 |
| | Feb | 16,040 | 16,950 | 910 | 6% | 0.292 |
| | Mar | 10,098 | 9,812 | -286 | -3% | 0.929 |
| | Apr | 5,325 | 5,543 | 218 | 4% | 0.790 |
| | May | 3,537 | 3,281 | -256 | -7% | 0.986 |
| | Jun | 2,679 | 2,682 | 3 | 0% | 0.817 |
| | Jul | 984 | 487 | -496 | -50% | 0.010 |
| | Aug | 115 | 94 | -21 | -18% | 0.606 |
| | Sep | 74 | 89 | 15 | 21% | 0.444 |
| | Oct | 158 | 222 | 64 | 41% | 0.028 |
| | Nov | 2,403 | 2,069 | -334 | -14% | 0.191 |
| | Dec | 5,970 | 6,382 | 412 | 7% | 0.958 |

Table S6: SONEC flooded agriculture - P1 (1988-2004) and P2 (2005-20) median monthly surface water change. Grass hay cultivation accounted for the vast majority of flooded agriculture, with other crops (e.g., wheat) making up a minor component of overall abundance.

| Month | P1 (1988-2004) | P2 (2005-2020) | Difference | % Difference | Wilcox *p* |
|---|---|---|---|---|---|
| Jan | 28,567 | 33,080 | 4,513 | 7% | 0.683 |
| Feb | 51,834 | 33,785 | -18,049 | -21% | 0.025 |
| Mar | 35,981 | 29,418 | -6,563 | -10% | 0.11 |
| Apr | 15,849 | 18,552 | 2,703 | 8% | 0.817 |
| May | 10,636 | 11,798 | 1,162 | 5% | 0.631 |
| Jun | 4,797 | 6,104 | 1,307 | 12% | 0.217 |
| Jul | 1,541 | 986 | -555 | -22% | 0.006 |
| Aug | 565 | 621 | 56 | 5% | 0.276 |
| Sep | 657 | 607 | -50 | -4% | 0.79 |
| Oct | 910 | 1,689 | 779 | 30% | 0.0579 |
| Nov | 5,474 | 6,717 | 1,243 | 10% | 0.444 |
| Dec | 15,364 | 20,311 | 4,947 | 14% | 0.276 |

Table S7: Central Valley all wetlands - P1 (1988-2004) and P2 (2005-20) median monthly surface water change. Areas (ha) include wetlands associated with duck clubs and wildlife refuges.

| Hydroperiod | Month | P1 (1988-2004) | P2 (2004-2020) | Difference | % Difference | Wilcox *p* |
|---|---|---|---|---|---|---|
| semi-permanent | Jan | 50,027 | 39,971 | -10,055 | -20% | 0.009 |
| | Feb | 48,900 | 42,706 | -6,195 | -13% | 0.015 |
| | Mar | 51,009 | 45,724 | -5,285 | -10% | 0.041 |
| | Apr | 46,319 | 41,968 | -4,351 | -9% | 0.001 |
| | May | 36,994 | 30,988 | -6,006 | -16% | 0.000 |
| | Jun | 28,293 | 25,424 | -2,870 | -10% | 0.000 |
| | Jul | 23,931 | 20,384 | -3,548 | -15% | 0.000 |
| | Aug | 22,419 | 19,726 | -2,692 | -12% | 0.000 |
| | Sep | 26,156 | 25,976 | -180 | -1% | 0.402 |
| | Oct | 40,891 | 38,760 | -2,131 | -5% | 0.127 |
| | Nov | 48,298 | 42,355 | -5,944 | -12% | 0.003 |
| | Dec | 44,996 | 38,328 | -6,668 | -15% | 0.008 |
| seasonal | Jan | 36,733 | 31,533 | -5,201 | -14% | 0.581 |
| | Feb | 45,825 | 33,946 | -11,879 | -26% | 0.136 |
| | Mar | 50,501 | 50,452 | -48 | 0% | 0.901 |
| | Apr | 38,922 | 29,354 | -9,568 | -25% | 0.000 |
| | May | 19,058 | 13,085 | -5,973 | -31% | 0.000 |
| | Jun | 8,003 | 5,592 | -2,412 | -30% | 0.000 |
| | Jul | 3,727 | 2,149 | -1,578 | -42% | 0.000 |
| | Aug | 2,496 | 1,654 | -842 | -34% | 0.000 |
| | Sep | 4,416 | 4,768 | 352 | 8% | 0.510 |
| | Oct | 16,910 | 19,477 | 2,567 | 15% | 0.001 |
| | Nov | 37,080 | 34,713 | -2,366 | -6% | 0.402 |
| | Dec | 42,304 | 41,050 | -1,253 | -3% | 0.465 |
| temporary | Jan | 15,661 | 10,588 | -5,073 | -32% | 0.683 |
| | Feb | 20,653 | 9,249 | -11,404 | -55% | 0.002 |
| | Mar | 17,209 | 27,987 | 10,779 | 63% | 0.345 |
| | Apr | 18,074 | 10,163 | -7,910 | -44% | 0.000 |
| | May | 10,296 | 5,896 | -4,400 | -43% | 0.657 |
| | Jun | 1,903 | 1,406 | -498 | -26% | 0.204 |
| | Jul | 886 | 401 | -485 | -55% | 0.002 |
| | Aug | 747 | 323 | -424 | -57% | 0.001 |
| | Sep | 1,253 | 920 | -333 | -27% | 0.008 |
| | Oct | 3,406 | 3,872 | 466 | 14% | 0.817 |
| | Nov | 10,179 | 13,151 | 2,972 | 29% | 0.094 |
| | Dec | 18,601 | 22,898 | 4,298 | 23% | 0.557 |

Table S8: Central Valley wildlife refuges - P1 (1988-2004) and P2 (2005-20) median monthly surface water change. Areas (ha) exclusive to state and federally managed wildlife refuges.

| Hydroperiod | Month | P1 (1988-2004) | P2 (2004-2020) | Difference | % Difference | Wilcox *p* |
|---|---|---|---|---|---|---|
| semi-permanent | Jan | 9,005 | 8,562 | -442 | -5% | 0.157 |
| | Feb | 9,370 | 8,602 | -768 | -8% | 0.015 |
| | Mar | 9,445 | 8,961 | -485 | -5% | 0.136 |
| | Apr | 8,987 | 8,312 | -674 | -8% | 0.006 |
| | May | 5,472 | 4,545 | -927 | -17% | 0.000 |
| | Jun | 3,801 | 3,150 | -651 | -17% | 0.000 |
| | Jul | 3,184 | 2,459 | -725 | -23% | 0.000 |
| | Aug | 2,921 | 2,364 | -558 | -19% | 0.000 |
| | Sep | 5,012 | 4,326 | -686 | -14% | 0.001 |
| | Oct | 7,879 | 7,678 | -201 | -3% | 0.157 |
| | Nov | 8,360 | 8,074 | -286 | -3% | 0.402 |
| | Dec | 7,797 | 7,521 | -276 | -4% | 0.423 |
| seasonal | Jan | 6,483 | 6,902 | 419 | 7% | 0.683 |
| | Feb | 6,830 | 7,086 | 256 | 4% | 0.709 |
| | Mar | 7,530 | 8,357 | 827 | 11% | 0.025 |
| | Apr | 5,158 | 4,866 | -292 | -6% | 0.231 |
| | May | 2,205 | 1,253 | -952 | -43% | 0.000 |
| | Jun | 1,002 | 482 | -519 | -52% | 0.000 |
| | Jul | 537 | 231 | -306 | -57% | 0.000 |
| | Aug | 284 | 156 | -129 | -45% | 0.000 |
| | Sep | 708 | 630 | -78 | -11% | 0.292 |
| | Oct | 2,840 | 3,257 | 417 | 15% | 0.049 |
| | Nov | 3,922 | 5,044 | 1,123 | 29% | 0.000 |
| | Dec | 3,833 | 5,213 | 1,380 | 36% | 0.000 |
| temporary | Jan | 2,854 | 1,895 | -959 | -34% | 0.041 |
| | Feb | 3,958 | 1,933 | -2,026 | -51% | 0.001 |
| | Mar | 3,976 | 5,136 | 1,159 | 29% | 0.345 |
| | Apr | 2,368 | 1,682 | -686 | -29% | 0.049 |
| | May | 645 | 355 | -289 | -45% | 0.000 |
| | Jun | 210 | 70 | -140 | -67% | 0.000 |
| | Jul | 90 | 37 | -53 | -59% | 0.000 |
| | Aug | 39 | 35 | -4 | -9% | 0.326 |
| | Sep | 128 | 148 | 20 | 16% | 0.423 |
| | Oct | 372 | 397 | 25 | 7% | 0.901 |
| | Nov | 758 | 562 | -196 | -26% | 0.345 |
| | Dec | 946 | 928 | -18 | -2% | 0.763 |

Table S9: Central Valley duck clubs - P1 (1988-2004) and P2 (2005-20) median monthly surface water change. Areas (ha) encompass private managed wetlands on duck clubs and wildlife preserves.

| Hydroperiod | Month | P1 (1988-2004) | P2 (2004-2020) | Difference | % Difference | Wilcox *p* |
|---|---|---|---|---|---|---|
| | Jan | 18,106 | 17,435 | -671 | -4% | 0.245 |
| | Feb | 19,614 | 18,062 | -1,552 | -8% | 0.049 |
| | Mar | 19,372 | 18,797 | -575 | -3% | 0.245 |
| | Apr | 17,655 | 16,108 | -1,547 | -9% | 0.009 |
| | May | 13,103 | 11,698 | -1,405 | -11% | 0.001 |
| semi-permanent | Jun | 10,305 | 8,883 | -1,422 | -14% | 0.000 |
| | Jul | 7,595 | 6,634 | -962 | -13% | 0.000 |
| | Aug | 6,965 | 6,161 | -804 | -12% | 0.000 |
| | Sep | 8,735 | 8,866 | 131 | 2% | 0.736 |
| | Oct | 15,700 | 16,090 | 391 | 2% | 0.245 |
| | Nov | 17,180 | 16,700 | -480 | -3% | 0.146 |
| | Dec | 16,479 | 15,163 | -1,316 | -8% | 0.157 |
| | Jan | 16,563 | 16,604 | 42 | >1% | 0.986 |
| | Feb | 17,315 | 17,163 | -152 | -1% | 0.901 |
| | Mar | 18,092 | 17,842 | -251 | -1% | 0.465 |
| | Apr | 10,673 | 8,728 | -1,945 | -18% | 0.000 |
| | May | 7,111 | 3,901 | -3,210 | -45% | 0.000 |
| seasonal | Jun | 3,905 | 1,552 | -2,353 | -60% | 0.000 |
| | Jul | 1,157 | 625 | -532 | -46% | 0.000 |
| | Aug | 567 | 425 | -142 | -25% | 0.004 |
| | Sep | 1,352 | 1,461 | 109 | 8% | 0.231 |
| | Oct | 7,915 | 9,886 | 1,971 | 25% | 0.000 |
| | Nov | 12,440 | 12,814 | 374 | 3% | 0.094 |
| | Dec | 11,641 | 13,842 | 2,201 | 19% | 0.037 |
| | Jan | 8,283 | 5,199 | -3,084 | -37% | 0.326 |
| | Feb | 8,744 | 4,226 | -4,518 | -52% | 0.001 |
| | Mar | 7,702 | 9,929 | 2,228 | 29% | 0.402 |
| | Apr | 4,124 | 2,816 | -1,308 | -32% | 0.007 |
| | May | 2,431 | 1,351 | -1,080 | -44% | 0.000 |
| temporary | Jun | 962 | 340 | -622 | -65% | 0.000 |
| | Jul | 269 | 165 | -104 | -39% | 0.002 |
| | Aug | 145 | 154 | 8 | 6% | 0.845 |
| | Sep | 309 | 242 | -67 | -22% | 0.041 |
| | Oct | 1,200 | 879 | -321 | -27% | 0.631 |
| | Nov | 2,775 | 2,167 | -607 | -22% | 0.309 |
| | Dec | 6,299 | 3,802 | -2,497 | -40% | 0.127 |

Table S10: Central Valley flooded agriculture - P1 (1988-2004) and P2 (2005-20) median monthly surface water change. Rice production accounted for the vast majority of flooded agriculture, with other crops (e.g., corn, wheat, and safflower) making up a relatively small component of overall abundance.

| Month | P1 (1988-2004) | P2 (2005-2020) | Difference | % Difference | Wilcox *p* |
|---|---|---|---|---|---|
| Jan | 100,562 | 129,178 | 28,616 | 29% | 0.094 |
| Feb | 126,728 | 107,588 | -19,140 | -15% | 0.168 |
| Mar | 111,633 | 114,714 | 3,081 | 3% | 0.901 |
| Apr | 102,683 | 71,447 | -31,235 | -30% | 0.000 |
| May | 187,093 | 200,724 | 13,631 | 7% | 0.118 |
| Jun | 92,598 | 107,864 | 15,266 | 17% | 0.053 |
| Jul | 7,764 | 6,685 | -1,079 | -14% | 0.110 |
| Aug | 2,450 | 2,141 | -308 | -13% | 0.069 |
| Sep | 6,306 | 3,716 | -2,590 | -41% | 0.002 |
| Oct | 33,758 | 22,747 | -11,011 | -33% | 0.002 |
| Nov | 62,253 | 109,335 | 47,082 | 76% | 0.002 |
| Dec | 70,650 | 118,618 | 47,969 | 68% | 0.001 |

Table S11: Major Reservoir storage in SONEC and the Central Valley (CV) - $km^3$ = cubic kilometers

| **Region** | **Minimum** | **1st Quartile** | **Median** | **Mean** | **3rd Quartile** | **Maximum** |
|---|---|---|---|---|---|---|
| SONEC (taf) | 110 | 455 | 648 | 654 | 837 | 1,223 |
| CV (taf) | 5,890 | 10,888 | 14,464 | 13,831 | 16,624 | 20,739 |
| CV/SONEC | 54 | 24 | 22 | 21 | 20 | 17 |
| SONEC ($km^3$) | 0.136 | 0.561 | 0.799 | 0.807 | 1.032 | 1.509 |
| CV ($km^3$) | 7.265 | 13.430 | 17.841 | 17.060 | 20.505 | 25.581 |

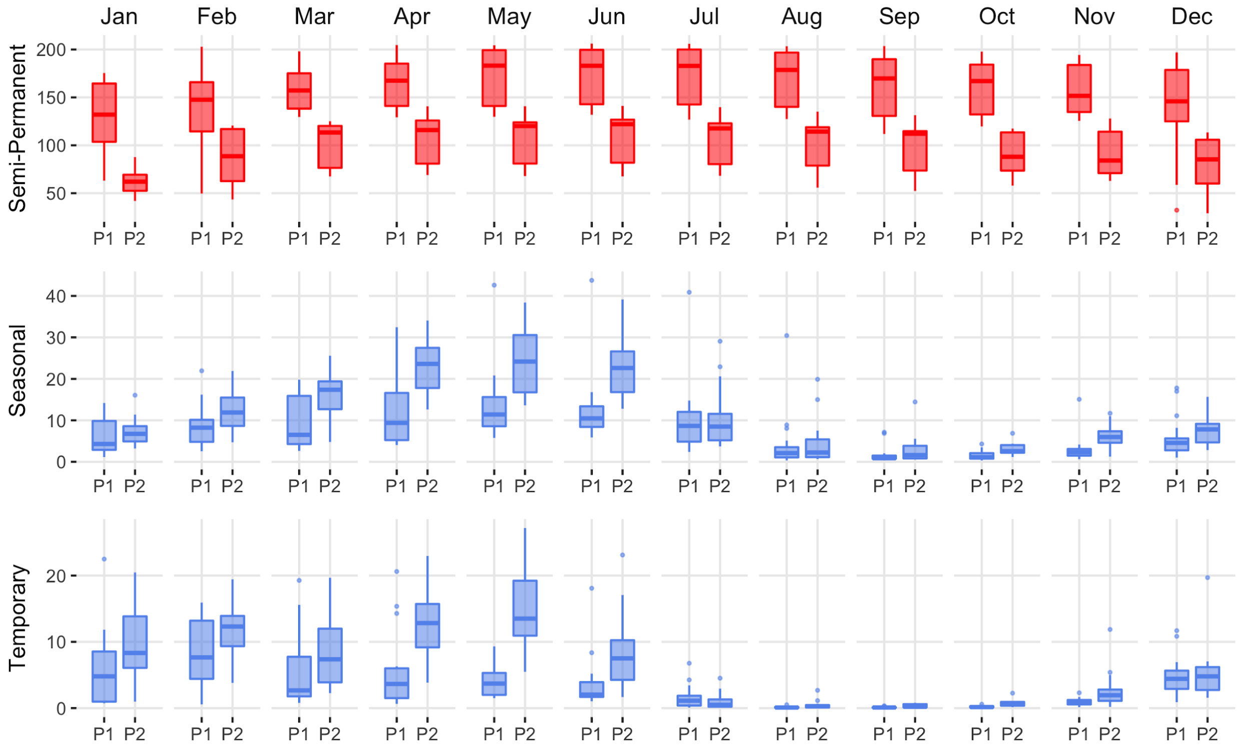


Figure S5. SONEC closed-basin lakes distribution of monthly wetland abundance (kha) between 1988-2004 (P1) and 2005-20 (P2) periods. Statistical inference determined as p-values < 0.1 derived from Wilcoxon ranked order test. Red indicates significant wetland decline and 'blue', stable to increasing wetland abundance. Results are partitioned by wetland hydroperiod (semi-permanent, seasonal, temporary). Boxes, interquartile range (IQR); line dividing the box horizontally, median value; whiskers, 1.5 times the IQR; points, outliers.

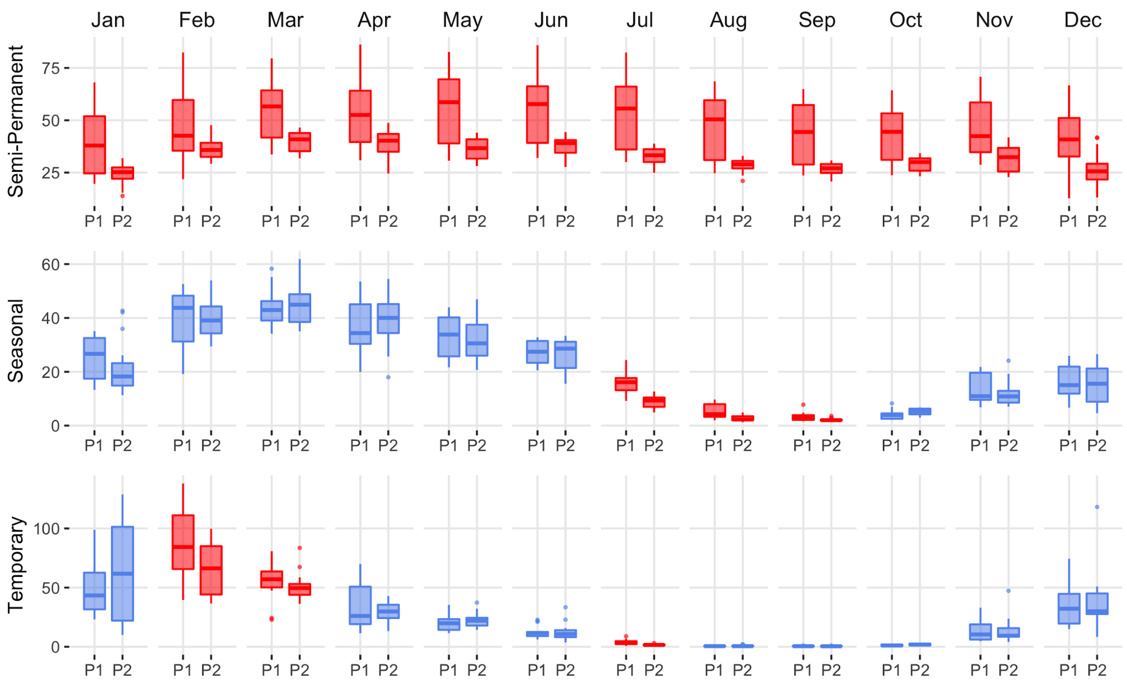


Figure S6: SONEC wildlife refuges - distribution of monthly wetland abundance (kha) from 1988-2004 (P1) and 2005-20 (P2). Areas exclusive to state and federally managed wildlife refuges. Statistical inference determined as p-values < 0.1 derived from Wilcoxon ranked order test. Red indicates significant wetland decline and ‘blue’, stable to expanding wetland abundance. Results are partitioned by wetland hydroperiod (semi-permanent, seasonal, temporary). Boxes, interquartile range (IQR); line dividing the box horizontally, median value; whiskers, 1.5 times the IQR; points, potential outliers.

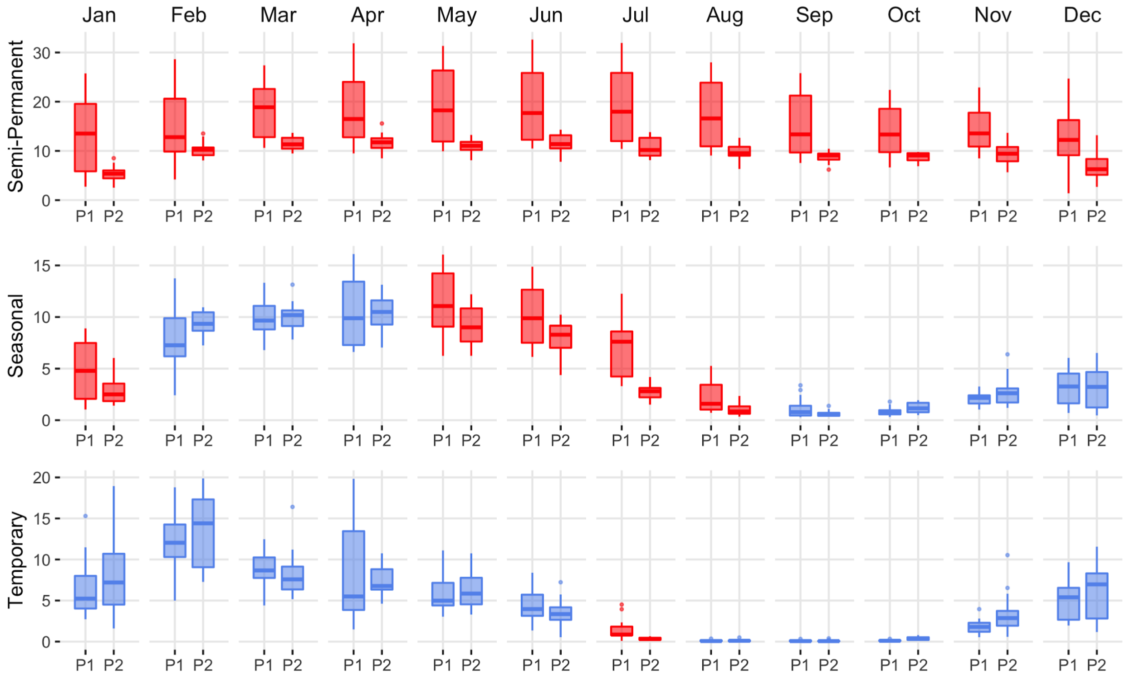


Figure S7: SONEC public wetlands - distribution of monthly wetland abundance (kha) from 1988-2004 (P1) and 2005-20 (P2) Areas include but are not limited to National Forest, Bureau of Land Management, and State Lands. Statistical inference determined as p-values < 0.1 derived from Wilcoxon ranked order test. Red indicates significant wetland decline and 'blue', stable to expanding wetland abundance. Results are partitioned by wetland hydroperiod (semi-permanent, seasonal, temporary). Boxes, interquartile range (IQR); line dividing the box horizontally, median value; whiskers, 1.5 times the IQR; points, potential outliers.

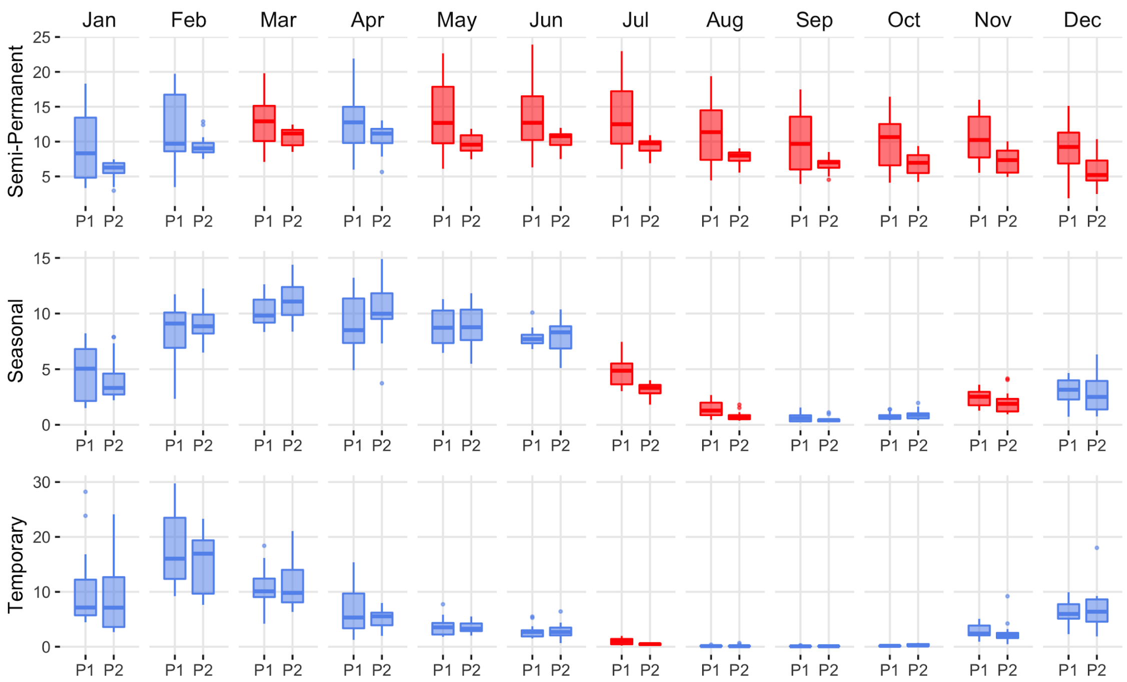


Figure S8: SONEC private wetlands - distribution of monthly wetland abundance (kha) from 1988-2004 (P1) and 2005-20 (P2). Areas were exclusive to wetlands on private lands not associated with agriculture. Statistical inference determined as p-values < 0.1 derived from Wilcoxon ranked order test. Red indicates significant wetland decline and ‘blue’, stable to expanding wetland abundance. Results are partitioned by wetland hydroperiod (semi-permanent, seasonal, temporary). Boxes, interquartile range (IQR); line dividing the box horizontally, median value; whiskers, 1.5 times the IQR; points, potential outliers.

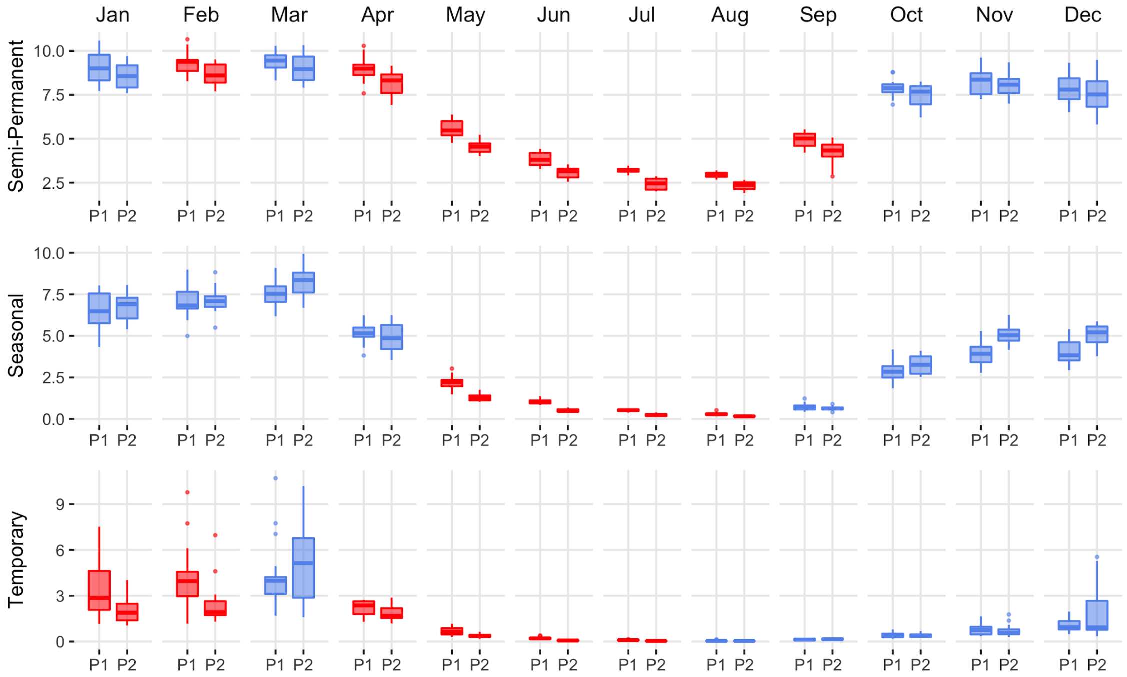


Figure S9: Central Valley wildlife refuges - distribution of monthly wetland abundance (kha) from 1988-2004 (P1) and 2005-20 (P2). Areas exclusive to state and federally managed wildlife refuges. Statistical inference determined as p-values < 0.1 derived from Wilcoxon ranked order test. Red indicates significant wetland decline and 'blue', stable to expanding wetland abundance. Results are partitioned by wetland hydroperiod (semi-permanent, seasonal, temporary). Boxes, interquartile range (IQR); line dividing the box horizontally, median value; whiskers, 1.5 times the IQR; points, potential outliers.

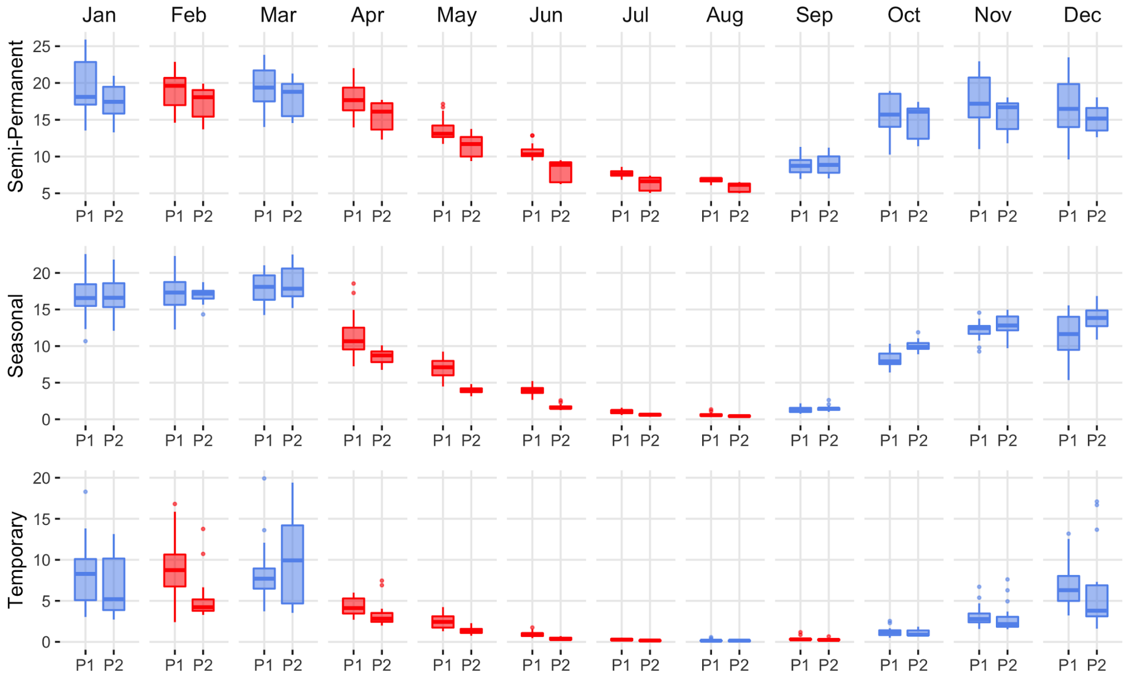


Figure S10: Central Valley duck clubs - distribution of monthly wetland abundance (kha) from 1988-2004 (P1) and 2005-20 (P2). Areas were exclusive to privately owned wetlands managed as waterfowl hunting preservers. Statistical inference determined as p-values < 0.1 derived from Wilcoxon ranked order test. Red indicates significant wetland decline and 'blue', stable to expanding wetland abundance. Results are partitioned by wetland hydroperiod (semi-permanent, seasonal, temporary). Boxes, interquartile range (IQR); line dividing the box horizontally, median value; whiskers, 1.5 times the IQR; points, potential outliers.

**Supplemental Materials – Recent Climate**

The regional water balance between source and use ultimately controls the surface water available for human use and wetland ecological systems in the SONEC and the Central Valley regions. The total amount of surface water is determined by precipitation, evapotranspiration, and resulting runoff, which drive groundwater recharge. These processes are highly modified by direct human factors, such as water withdrawal for industrial, domestic, and agricultural use (AghaKouchak, et al., 2021). To examine climate change over the period of our analysis, we used the TerraClimate dataset (Abatzoglou *et al.* 2018), a gridded (4km) monthly climate and water balance model of terrestrial surfaces available through the Google Earth Engine platform (Gorelick, et al., 2017). TerraClimate data is compiled using a climatically aided interpolation of relatively high resolution spatial and temporal scales, which has been validated with data from a broad climate network, including evapotranspiration and runoff, important for determining hydro-climate change (http://www.climatologylab.org/terraclimate.html). Each month was smoothed with a 5-year rolling mean to match the approach used to calculate surface water estimates to minimize inter-annual variability from exogenous and endogenous drivers. Trends were compiled using periods aligned with surface water summaries (P1=1988-2004; P2=2005-2020) and compared using nonparametric Wilcoxon rank order tests (Siegel 1957). By comparing trends over long periods, we were able to minimize the effects of shorter-term climate cycles (e.g., El Nino Southern Oscillation; Dettinger *et al.* 1998) that may have influenced results. Overall results were provided as boxplots partitioned by climate variable and region. Data are presented in the following figures (Figures S5-S8) to show changes in climate variables over the periods of interest presented in the Results and Discussion sections. A p-value of $< 0.1$ was used to represent statistical significance. In plots, significance levels between periods are shown by symbols representing significant cut points:

**** $p < 0.0001$
*** $p < 0.001$
** $p < 0.01$
* $p < 0.1$.

Figure S11

**SONEC**

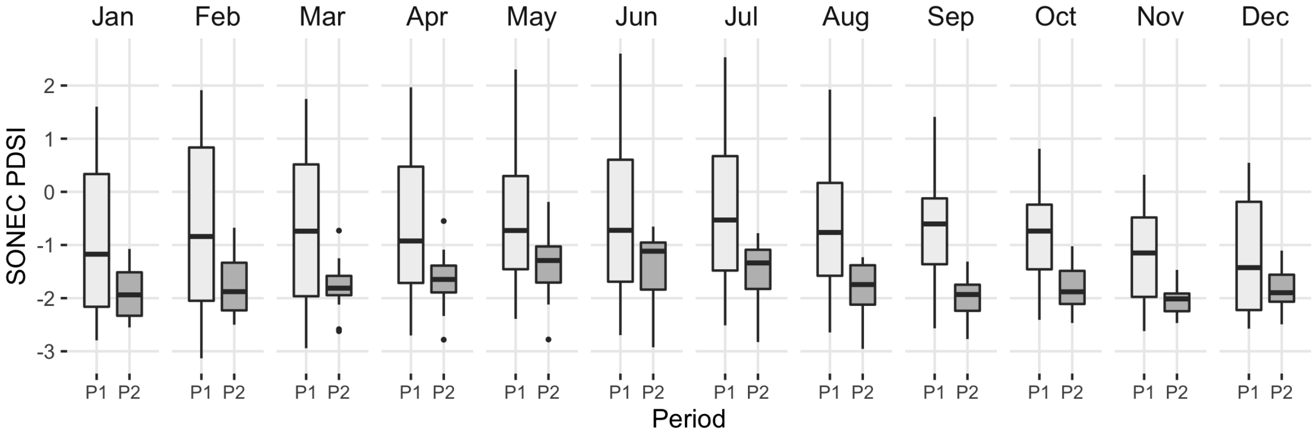


**Central Valley**

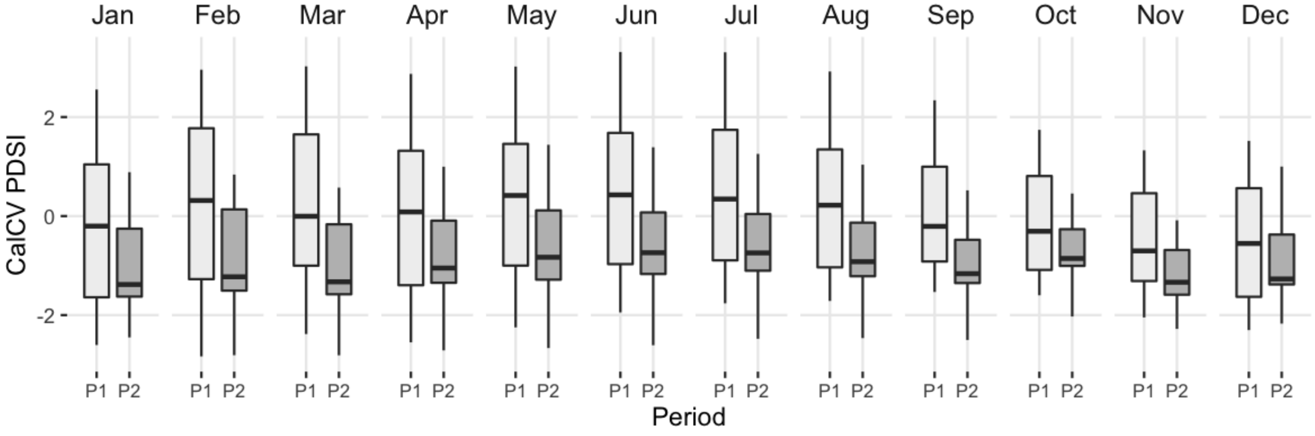

Figure S12

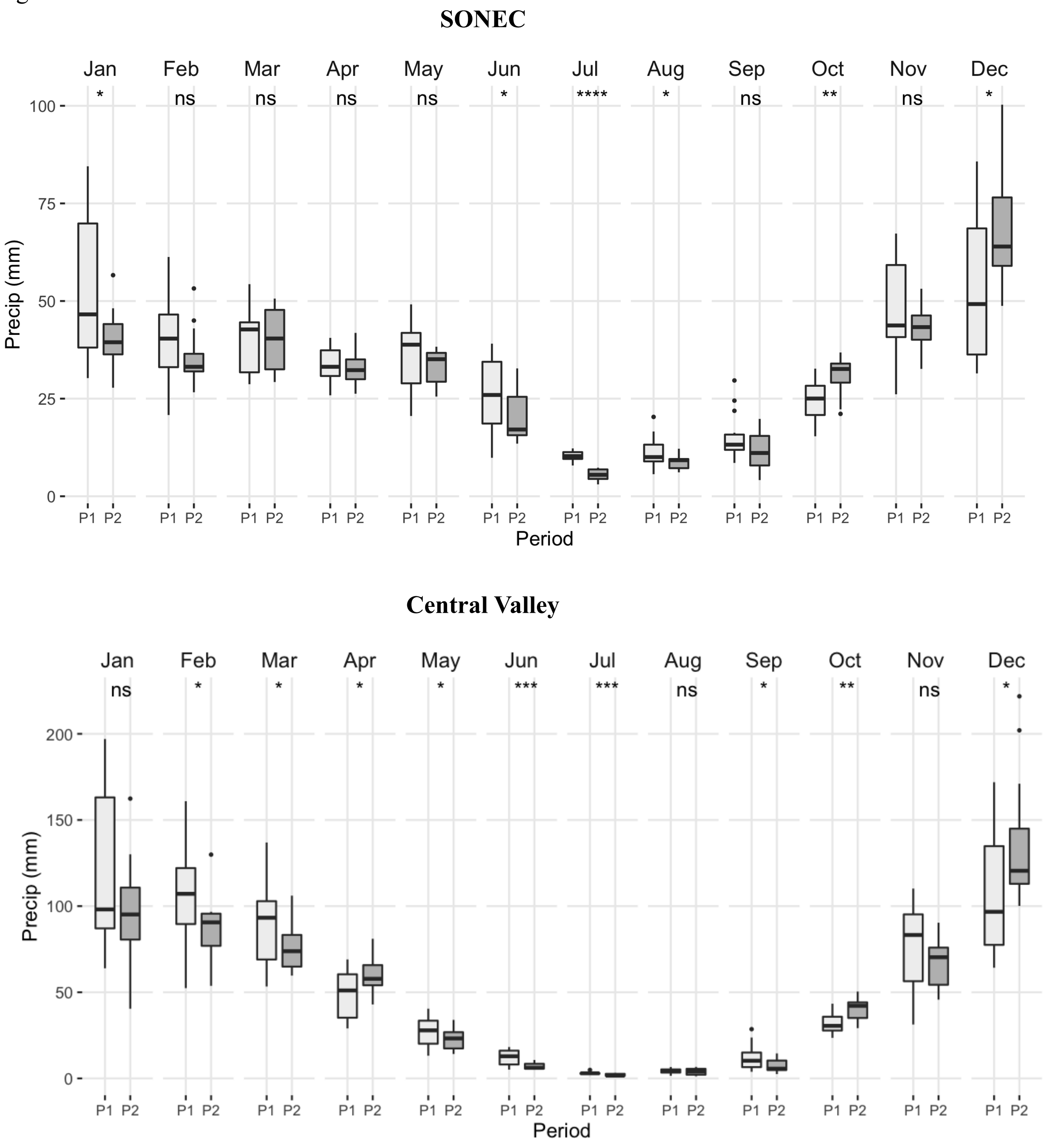
SONEC
Jan
Feb
Mar
Apr
May
Jun
Jul
Aug
Sep
Oct
Nov
Dec
* ns ns ns ns * **** * ns ** ns *
Precip (mm)
0
25
50
75
100
P1 P2
Period
Central Valley
Jan
Feb
Mar
Apr
May
Jun
Jul
Aug
Sep
Oct
Nov
Dec
ns * * * * *** *** ns * ** ns *
Precip (mm)
0
50
100
150
200
P1 P2
Period

Figure S13

**SONEC**

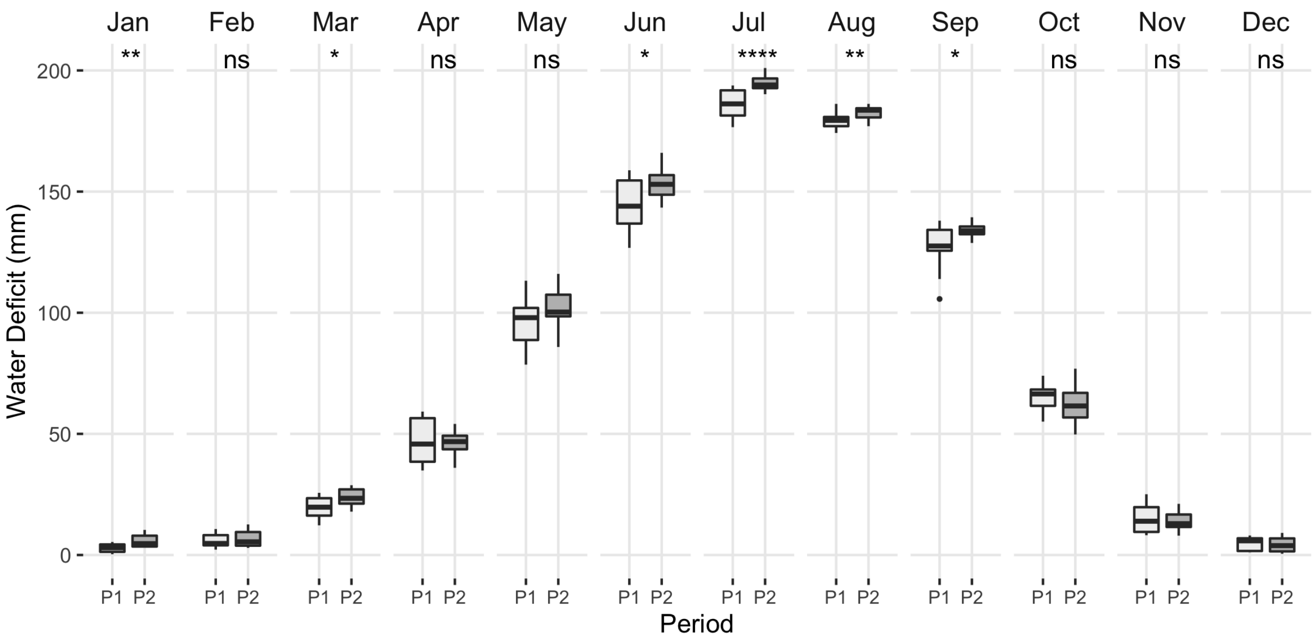


**Central Valley**

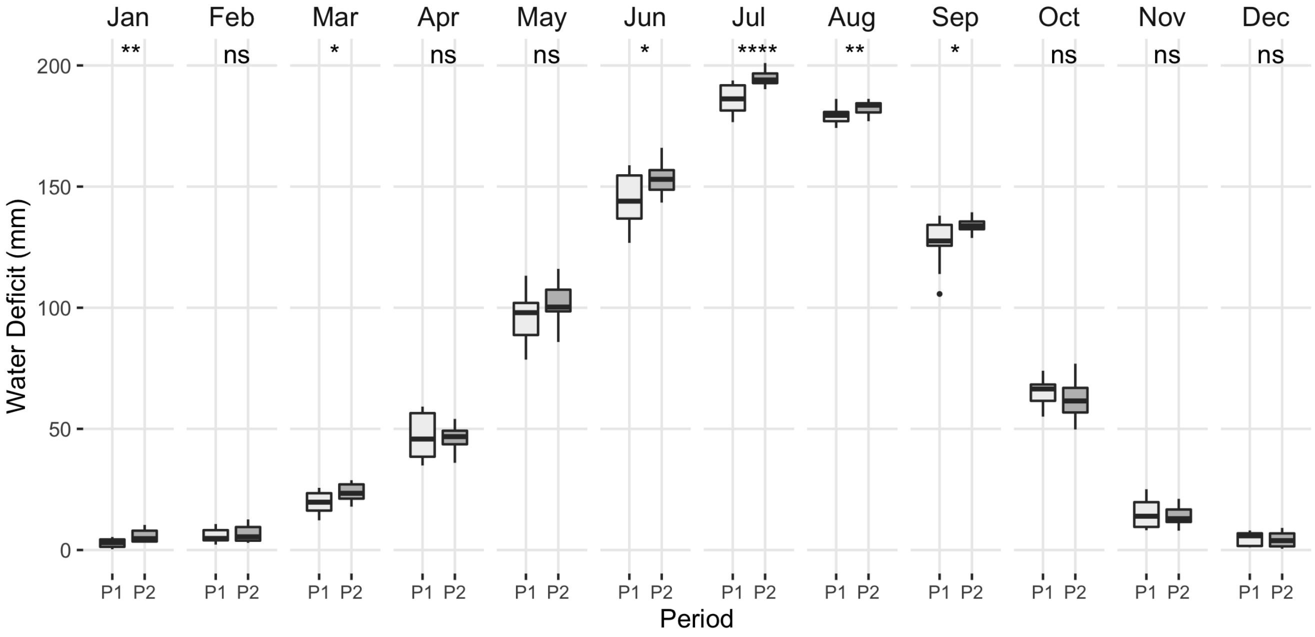

Figure S14

**SONEC**

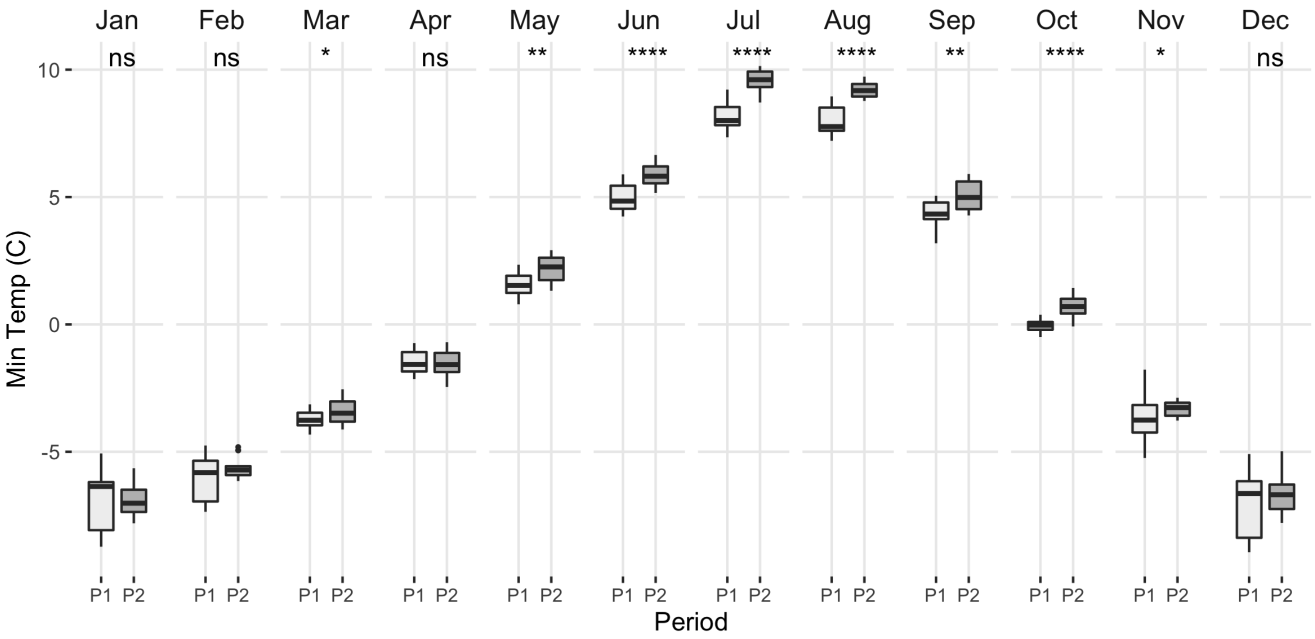


**Central Valley**

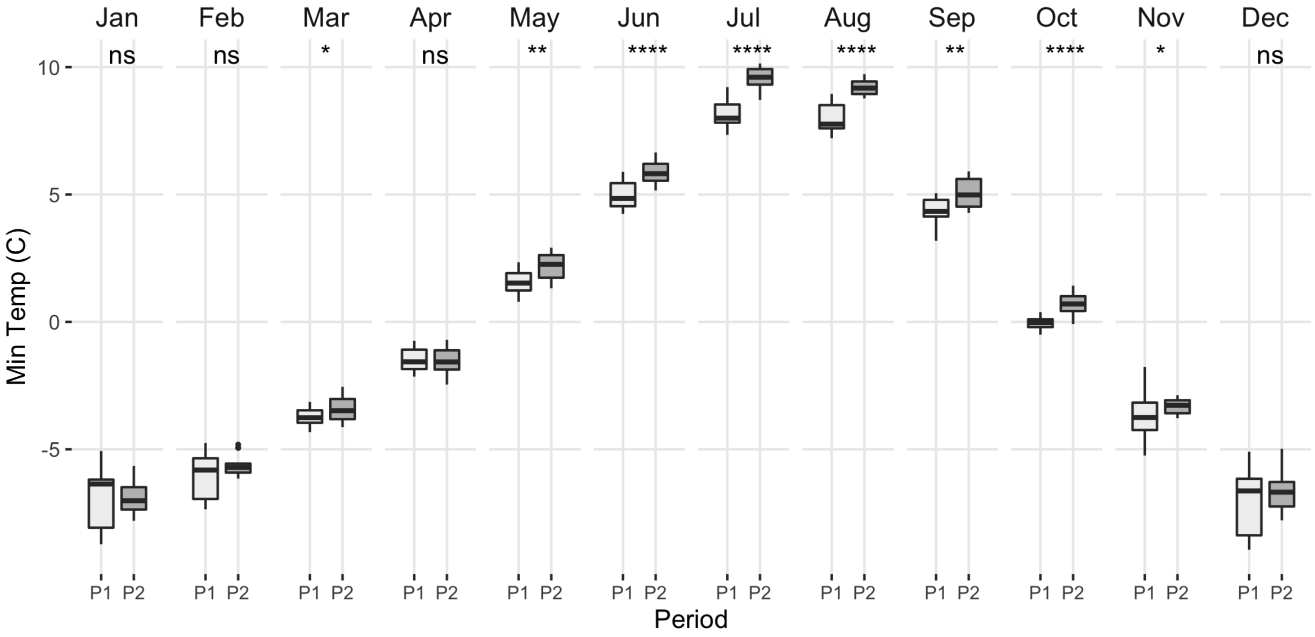

Figure S15

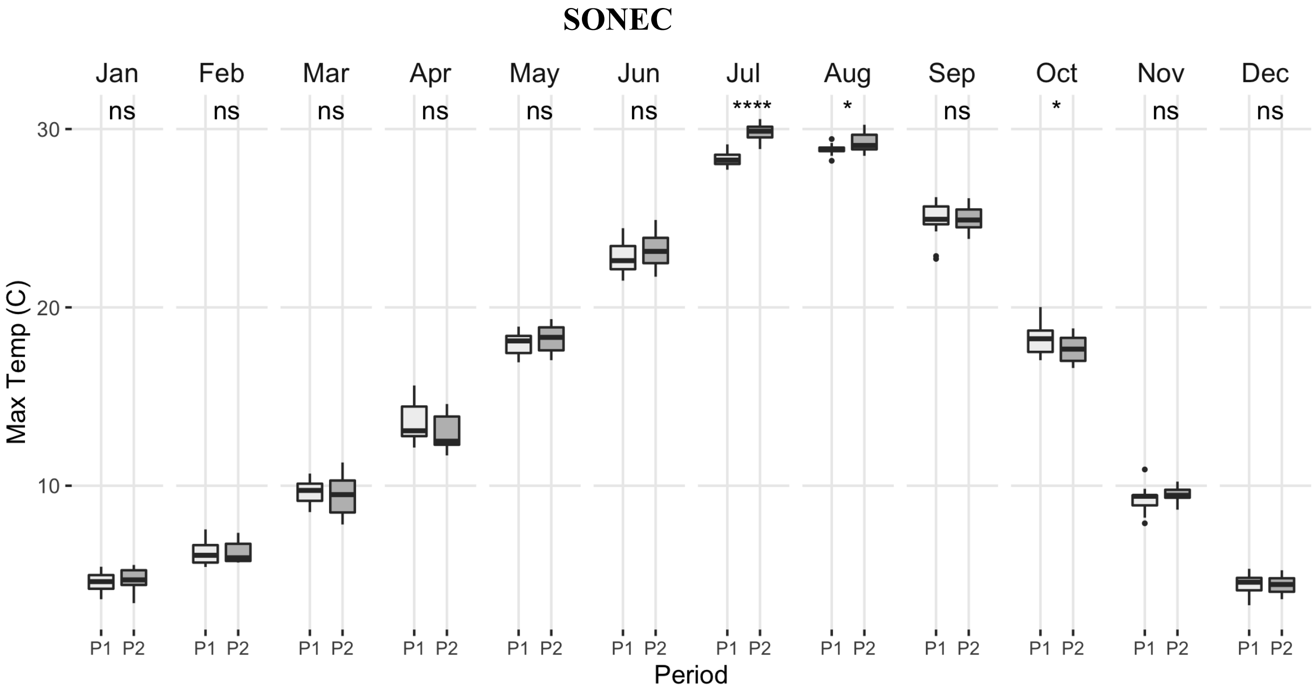
SONEC
Jan
Feb
Mar
Apr
May
Jun
Jul
Aug
Sep
Oct
Nov
Dec
ns
ns
ns
ns
ns
ns
****
*
ns
*
ns
ns
30
20
10
Max Temp (C)
P1 P2
Period

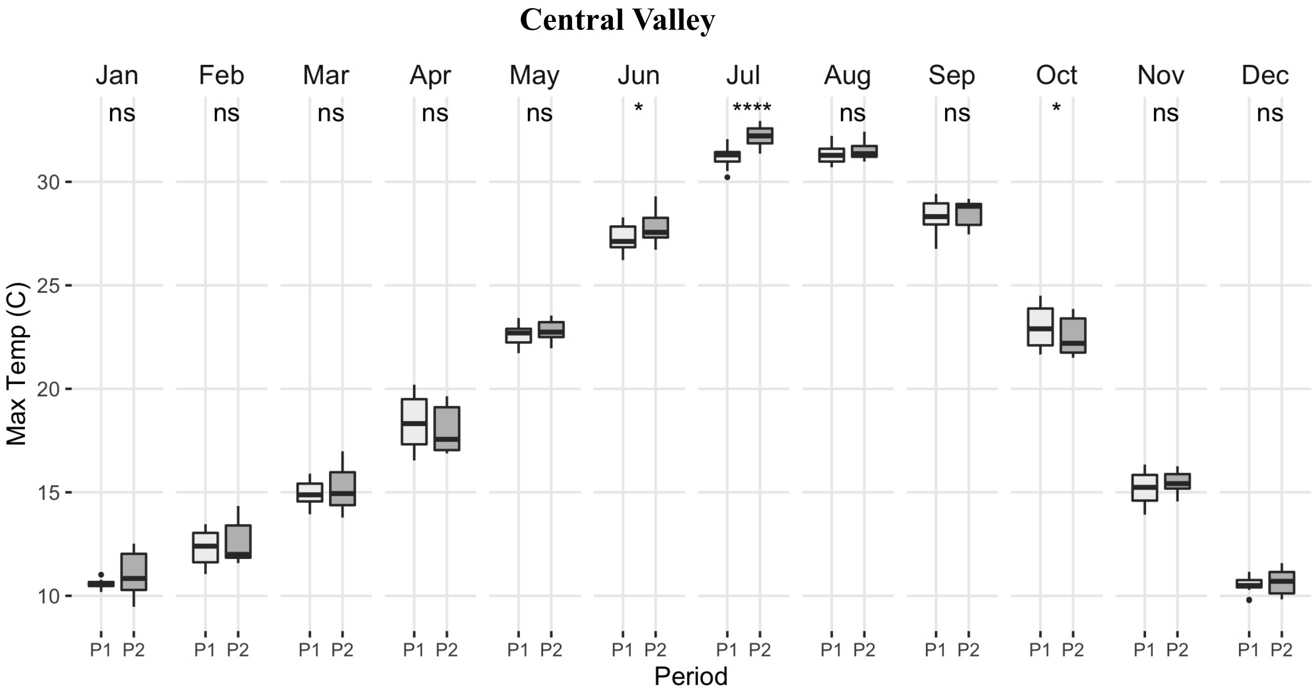
Central Valley
Jan
Feb
Mar
Apr
May
Jun
Jul
Aug
Sep
Oct
Nov
Dec
ns
ns
ns
ns
ns
*
****
ns
ns
*
ns
ns
30
25
20
15
10
Max Temp (C)
P1 P2
Period

**Supplemental Materials – Future Climate**

To examine the potential future climate of the SONEC and the Central Valley regions, we used the MACAv2-METDATA Monthly Summaries accessed in the Google Earth Engine Platform (GEE). The dataset is statistically downscaled from global climate model output from the Coupled Model Intercomparison Project 5 (CMIP5, Taylor et al. 2010) utilizing a modification of the Multivariate Adaptive Constructed Analogs approach (hence, MACA, Abatzoglou, and Brown, 2012) with the "Livneh" observational dataset as training data (Livneh et. al., 2013). MACAAv2 has a 4-km grid-scale for historical (1950-2005) and future (2006-2100) climate metrics (maximum and minimum temperature, humidity, precipitation, and downward shortwave radiation), compiled for 20 global climate models. Detailed information on methods and available data can be found at the MACA homepage (https://climate.northwestknowledge.net/MACA/index.php).

We used an ensemble of downscaled climate models to estimate variability in future climate outcomes (Mote, et al. 2011). To access both model and scenario uncertainty (Hawkins and Sutton 2009) we compiled data from a set of models run under two future representative concentration pathways (RCP) RCP 4.5 and RCP 8.5. RCP 4.5, an intermediate scenario, has $CO_2$-equivalent emissions peaking ~2040, then declining through 2100 (Fig. 2 in Meinshausen, et al. 2011; https://ar5-syr.ipcc.ch/topic_futurechanges.php, Box 2.2, Fig. 1). RCP 8.5 assumes emissions steadily rise through 2100 and, although "increasingly implausible with each passing year" (Hausfather and Peters 2020), represents a high-emission boundary condition or "worst-case" climate change scenario (*ibid*). RCP 8.5 and RCP4.5 temperature projections are approximately consistent with the model outcomes from the 2000 Special Report on Emission Scenarios (SRES) A1FI and B1, respectively (Hayhoe, et al., 2017, Fig. 4.1, p. 138), representing the potential global maximum temperature response (~ 4-10°C by 2100) and a more moderate response ~ 2-4°C by 2100).

Although some studies suggest that projections from a random set of climate models are similar to those of the "best" models based on comparison to historical data, we used results from Rupp et al. (2013) to inform model selection rather than using all 20 MACAv2 models. Rupp et al. (2013) found that for the Pacific Northwest region (which overlaps most of the SONEC and the

Central Valley regions), there was a significant difference among 41 downscaled climate models. However, a clear set of models did a better job of reproducing historical conditions over 18 climate metrics. This was especially true for the metrics measuring the seasonal amplitude and inter-annual/seasonal variability of precipitation and temperature, metrics important for understanding wetland response to climate and waterbird use of wetland systems. Therefore we decided to use the models that overlapped between the top-20 models of Rupp et al. and those available in MACAAv2. This resulted in a set of 7 models (Table S2).

***Table S11****: List of models used from the MACAv2 GEE dataset that are in the "best" performing models (top ~15) of Rupp, et al. (2013).*

| MODEL | SOURCE AGENCY |
|---|---|
| CanESM2 | Canadian Centre for Climate Modeling and Analysis |
| CCSM4 | National Center of Atmospheric Research, USA |
| HadGEM2‑CC | Met Office Hadley Center, UK |
| HadGEM2‑ES | Met Office Hadley Center, UK |
| IPSL-CM5B-MR | Institute Pierre Simon Laplace, France |
| MICROC5 | Japan (three institutes) |
| NorESM1-M | Norwegian Climate Center, Norway |

Although this is a relatively small number of models for ensemble analysis (https://climate.northwestknowledge.net/MACA/GCMselection.php), we chose this approach because of the lower relative error of variables of interest in this set of MACAv2-available model outcomes provides a stronger gage of variability/uncertainty of future climate projections in the SONEC-Central Valley regions.

We extracted MACAv2 values from these seven models for the RCP 4.5 and RCP 8.5 scenarios to determine precipitation and temperature (maximum and minimum) for each water year from the historical period from 1950-1999 and the future from 2039-2099 (three ~ 20-year periods). Rather than using modeled actual values, we calculated and plotted future anomalies for temperature and precipitation based on the 1950-1999 period median values (see following Figures S13 & S14). The changes in three climate variables, minimum temperature (TMIN), maximum temperature (TMAX), and precipitation (PR) in the future from this assemblage of models are presented. In all cases, anomalies are plotted based on the historical 1950-1999 period.

Figure S16: Future climate projections for SONEC

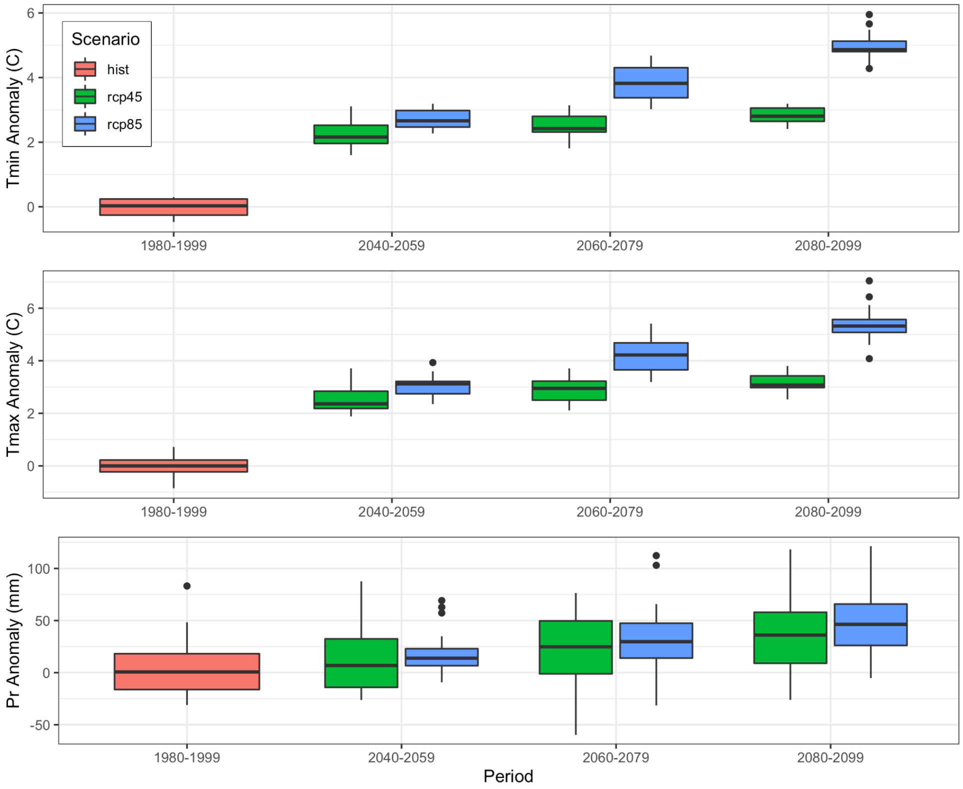

Figure S17: Future climate projections for Central Valley

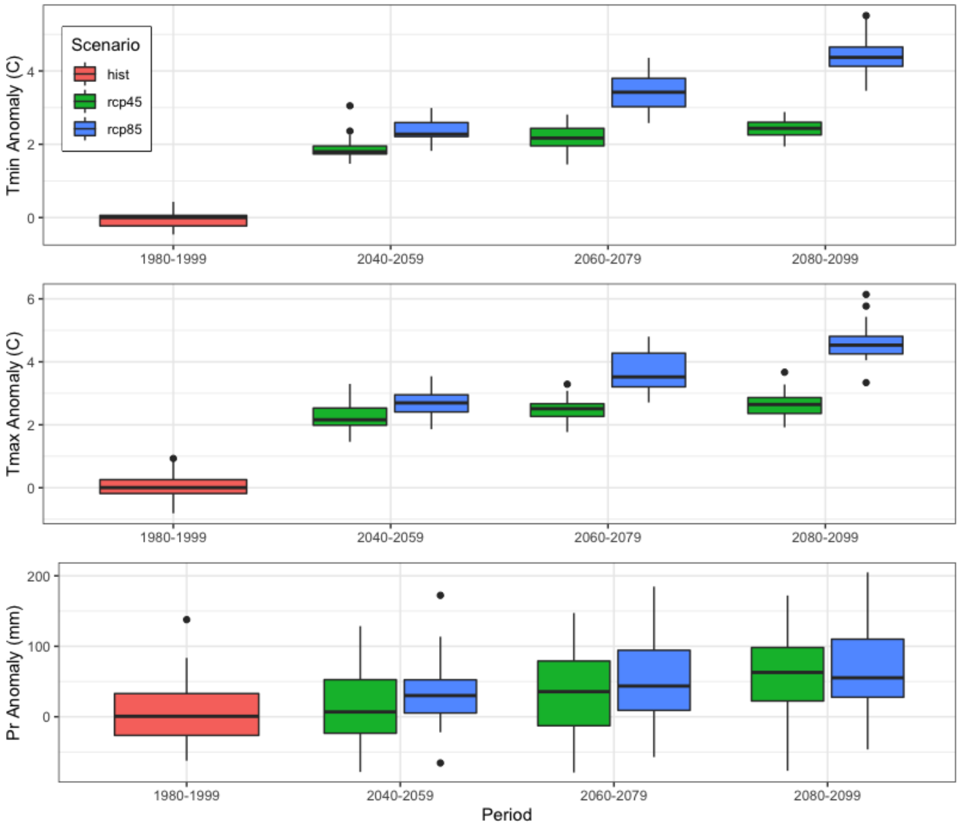


We also used MACAv2 values to calculate time series of the Standardized Precipitation Evapotranspiration Index (SPEI) (Vicente-Serrano, Sergio M. & National Center for Atmospheric Research Staff (Eds). Last modified 18 Jul 2015. "The Climate Data Guide: Standardized Precipitation Evapotranspiration Index (SPEI)." Retrieved from https://climatedataguide.ucar.edu/climate-data/standardized-precipitation-evapotranspiration-index-spei):

The SPEI considers precipitation (PRCP) and potential evapotranspiration (PET) in estimating drought, capturing the impact of temperature on water demand. The SPEI also correlates well with the self-calibrating Palmer Drought Severity Index (scPDS I) at 1-18 month timescales, the scale at which we examine changing wetland surface areas. To estimate potential drought in the future, we used MACAv2 data as input to the R-package SPEI (S.M. Vicente-Serrano, S. Beguería, J.I. López-Moreno. 2010. A Multi-scalar drought index sensitive to global warming: The Standardized Precipitation Evapotranspiration Index – SPEI. Journal of Climate 23: 1696, DOI: 10.1175/2009JCLI2909.1: URL http://sac.csic.es/spei). We calculated PET by the Hargreaves method in the SPEI package and then calculated the SPEI time series from 1950 to 2100, the time range of the MACAv2 data. We used the period just before our surface water analysis, 1950-1985, as the reference period. SPEI was then calculated for the complete time series using "historical", "rcp45", and "rcp85" scenarios in the MACAv2 data for each region, SONEC, and the Central Valley. This resulted in four time series representing potential future drought under an intermediate $CO_2$-equivalent emissions scenario (RCP 4.5) and a high-emission change scenario (RCP 8.5), presented in Figures S16 & S17.

Figure S18: SPEI for 1950-2100 for the SONEC region for rcp 4.5 and rcp 8.5 emissions scenarios. Reference period is shaded, 1950 -1985.

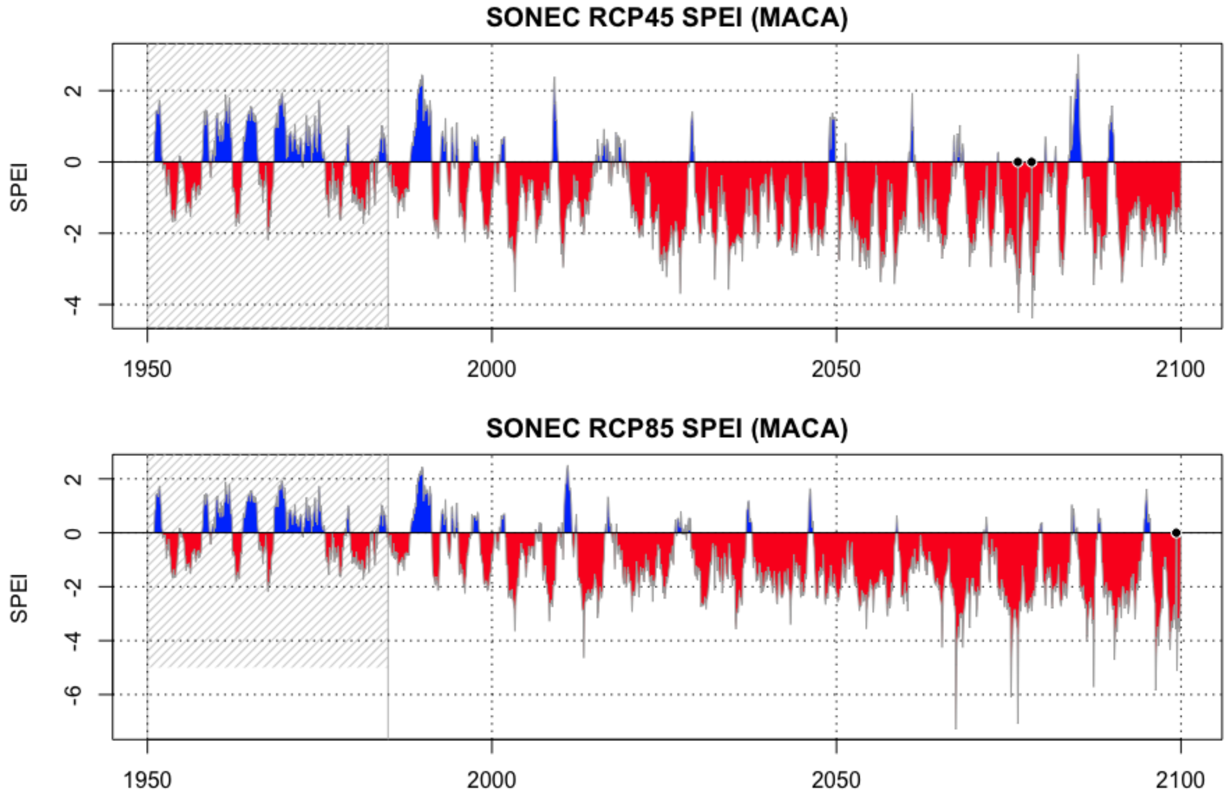

Figure S19: SPEI for 1950-2100 for the Central Valley region for rcp 4.5 and rcp 8.5 emissions scenarios. Reference period is shaded, 1950 -1985.

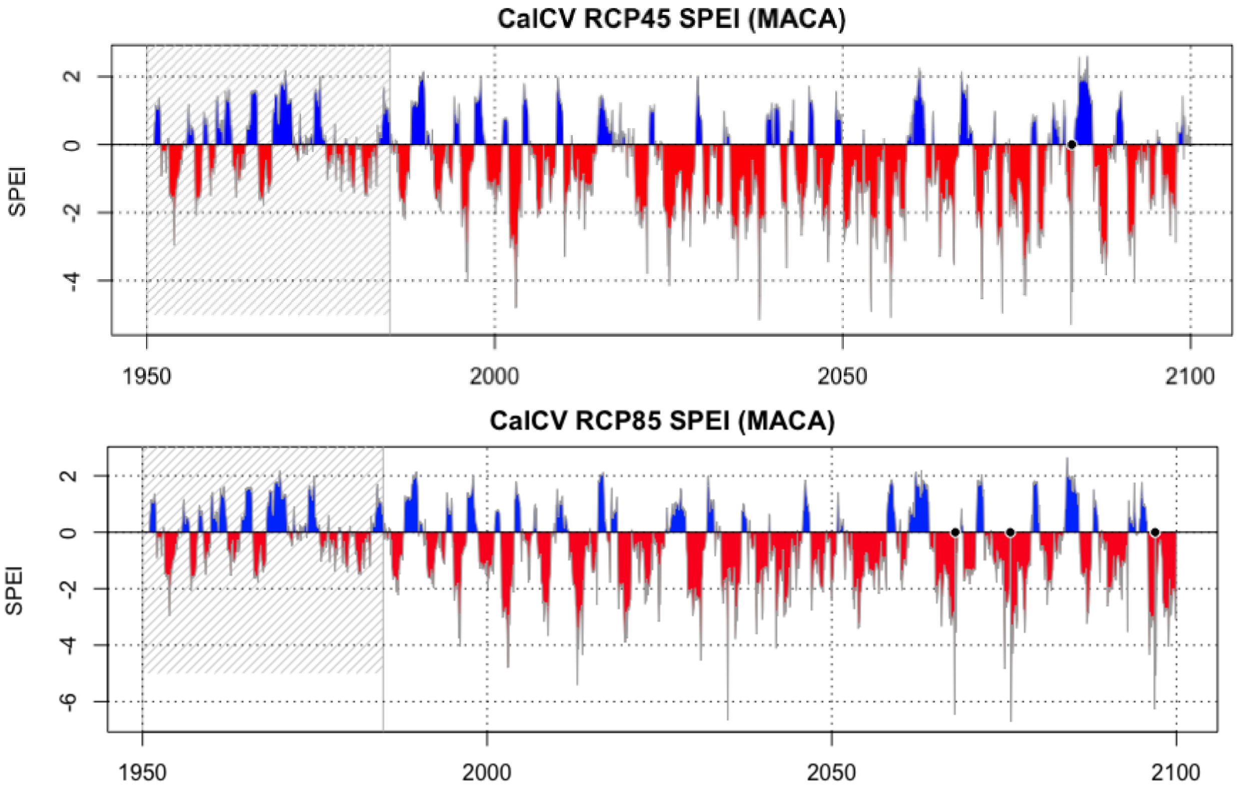
